# Oxytocin modulates neurocomputational mechanisms underlying prosocial reinforcement learning

**DOI:** 10.1101/2021.05.26.445739

**Authors:** Daniel Martins, Patricia Lockwood, Jo Cutler, Rosalyn Moran, Yannis Paloyelis

## Abstract

Humans often act in the best interests of others. However, how we learn which actions result in good outcomes for other people and the neurochemical systems that support this ‘prosocial learning’ remain poorly understood. Using computational models of reinforcement learning, functional magnetic resonance imaging and dynamic causal modelling, we examined how different doses of intranasal oxytocin, a neuropeptide linked to social cognition, impact how people learn to benefit others (prosocial learning) and whether this influence could be dissociated from how we learn to benefit ourselves (self-oriented learning). We show that a low dose of oxytocin prevented decreases in prosocial performance over time, despite no impact on self-oriented learning. Critically, oxytocin produced dose-dependent changes in the encoding of prediction errors (PE) in the midbrain-subgenual anterior cingulate cortex (sgACC) pathway specifically during prosocial learning. Our findings reveal a new role of oxytocin in prosocial learning by modulating computations of PEs in the midbrain-sgACC pathway.

## Introduction

Prosocial behaviours - actions intended to benefit other people - are crucial for social cohesion^1^. From small acts of kindness to major sacrifices, prosocial behaviours have intrigued many disciplines for centuries^2^. While debate persists about the intrinsic motives that guide us towards behaving prosocially, there is consensus that, in order to help, we must be able to learn the impact our actions have on others^2, 3^.

Reinforcement learning (RL) theory provides a neurobiologically plausible framework to explain how humans and other species form action-outcome associations^4^. Recent evidence has shown that humans rely on the same reinforcement learning algorithms when learning to benefit themselves (self-oriented learning)^3^ and others (prosocial learning). Yet these algorithms are implemented by distinct circuits in the brain and have different influences on behaviour^5^. Both self-oriented and prosocial reinforcement learning are driven by the difference between expected and actual outcomes, known as prediction errors (PE)^3^. PE are signalled through changes in the phasic release of dopamine in the forebrain^6, 7^ and drive learning by updating the expected value of future choice options^8, 9^. Humans learn faster when they are the recipients of the rewards themselves as compared to others (self-bias). The encoding of PE for prosocial and self-directed outcomes partially map to common anatomical substrates, such as the nucleus accumbens^3^. However, the encoding of prosocial PE specifically engages additional brain pathways anchored in the subgenual anterior cingulate cortex (sgACC)^3^, a region that is thought to play a key role in many aspects of social cognition^10^.

In addition to identifying the neuroanatomical pathways where prosocial learning computations take place, understanding the neurochemical systems that support prosocial learning and govern the neurocomputational mechanisms through which they are implemented is critical. Ultimately, this would allow us to identify putative molecular targets that could enhance prosocial behaviour in behavioural disorders characterised by dysfunctional social behaviour, such as antisocial behaviour, where we currently lack efficient therapies^11^.

Oxytocin, a hypothalamic neuropeptide repeatedly implicated in social cognition and behaviour^12^, is a strong molecular candidate for targeting prosocial learning and its underlying neurocomputational mechanisms. First, oxytocin plays a crucial role in the encoding of social feedback during learning, through interactions with the dopaminergic mesolimbic pathways^13^. Second, a single dose of intranasal oxytocin has been shown to modulate the neurocomputational processes that take place during reinforcement learning, i.e. intranasal oxytocin increases representations of social value in the amygdala during economical exchanges^14^ and blunts the encoding of PE when humans have to learn that others should not be trusted^15^. Third, the sgACC, where PEs are encoded during prosocial learning specifically, receives oxytocinergic innervation^16^ and expresses mRNA of the oxytocin receptor gene abundantly^17^. Taken together, these lines of evidence converge on the hypothesis that oxytocin might act as a biological facilitator of prosocial learning by impacting on the neural computations that take place in the midbrain-sgACC pathway when we learn to benefit others.

Here, we set out to test this hypothesis by examining the effect of three doses of intranasal oxytocin or placebo on self-oriented versus prosocial learning. We recruited 24 healthy men to participate in a double-blind, placebo-controlled, within-subjects, dose-response study where we administered 9, 18, 36 IU of intranasal oxytocin or placebo to each participant in four different days using a nebuliser (Figure 1a). We asked participants to perform a reinforcement learning task that can dissociate neural mechanisms for prosocial and self-benefitting learning^3^. In this task rewards would be paid either to the participant (self-oriented learning condition) or to a stranger (a confederate; prosocial learning condition), whom the participants was introduced to at the start of the study. On each trial participants had to choose between one of two abstract symbols. One symbol was associated with a high probability (75%) and one was associated with a low probability (25%) of obtaining a reward. These contingencies were not instructed but had to be learned through trial and error from feedback on whether the reward was received presented at the end of each trial (Figure 1b).

**Figure 1.**
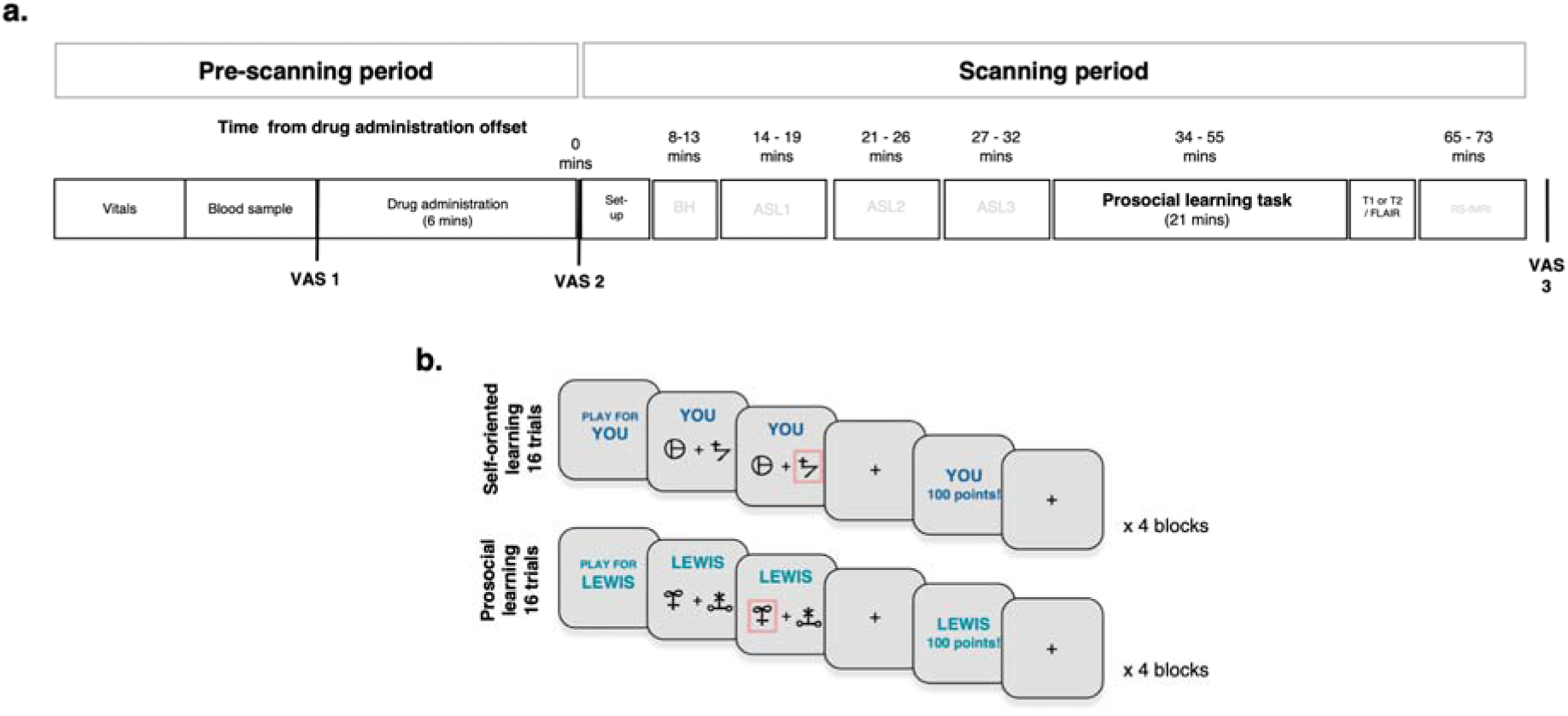
Protocol of the study (a) and prosocial reinforcement learning task (b). In panel (a), we provide an overview of the experimental procedures of our study. <u>Pre-Scanning period:</u> Each session started with a quick assessment of vitals (heart rate and blood pressure) and collection of two blood samples for plasma isolation. Then, participants self-administered one of three possible doses of intranasal oxytocin (∼9, 18 or 36 IU) or placebo using the PARI SINUS nebulizer. The participants used the nebulizer for three mins in each nostril (total administration 6 mins). Immediately before and after drug administration, participants filled a battery of visual analog scales (VAS) to assess subjective drug effects (alertness, mood and anxiety). <u>Scanning Period:</u> Participants were then guided to a magnetic resonance imaging scanner, where acquired BOLD-fMRI during a breath hold (BH) task, three consecutive arterial spin labelling (ASL) scans, the BOLD-fMRI prosocial learning task, followed by structural scans (T1 or T2 / FLAIR) and one resting-state fMRI (RS-fMRI) at the end. We present the time-interval post-dosing (mean time from drug administration offset) during which each scan took place. At the end of the scanning session, we repeated the same battery of VAS to subjective drug effects. In panel (b), we present an overview of the prosocial reinforcement learning task. Participants had to learn the probability that abstract symbols were rewarded to gain points over 16 trials in each block. At the beginning of each block, participants were told who they were playing for either themselves or for other participant (unbeknown to the participants, this other participant was a confederate). Points from the ‘self-oriented learning’ condition were converted into additional payment for the participant themselves, points from the ‘prosocial learning’ condition were converted into money for the other participant. Participants played four blocks in each condition.

Using computational models of reinforcement learning and functional magnetic resonance imaging (fMRI), we show that prosocial and self-orientated learning processes exhibit two key differences. First, participants are better at learning how to get rewards for themselves than for others (self-bias). Second, while performance for oneself is maintained at high levels throughout the task, performance declines over time when rewards are for someone else. Intranasal oxytocin produced a dose-response effect specific to the prosocial condition. Compared to placebo, a low dose of oxytocin, but not the medium or high doses, prevented the decrease in prosocial performance over time with no effect on self-orientated learning. Moreover, intranasal oxytocin produced dose-dependent changes in the encoding of PE in the midbrain-sgACC pathway during prosocial learning. A low dose, compared to placebo, strengthened the encoding of PE in this pathway by increasing excitatory midbrain-to-sgACC transmission, while a high dose decreased excitatory midbrain-to-sgACC transmission. Overall, we reveal both behavioural and neural influences of oxytocin on prosocial learning that can be dissociated from oxytocin’s effects on self-oriented learning. Our findings could have important implications for strategies to foster prosocial behaviours in health and disorder.

## Results

We confirmed that all participants believed our cover story at the end of their participation. Even though the prior contact between participants and confederates was standardized and kept to a minimum, humans form quick and strong first impressions about others^18^ which could then influence how prosocial learning evolved in the task. For this reason, we assessed participants’ impression of the confederate right after they interacted using an impression scale^19^. We then examined whether our confederates might have elicited any form of strong preference bias. We did not detect any significant differences between the average ratings of the confederates and the middle point of the impression scale (T(23) = -0.549, p = 0.588), which suggests that the confederates were perceived neutrally.

### Only a low dose of intranasal oxytocin prevents decreases in prosocial performance over time but does not impact on self-oriented learning

We first examined participants’ ability to complete the task in both learning conditions (self-oriented or prosocial) and all treatment levels (placebo, low, medium, high). Participants selected the option with the higher chance of receiving a reward significantly above chance (50%) during both self-oriented and prosocial learning in all treatment levels (smallest T(23) = 5.191, p = 0.007) (Supplementary Figure S2).

To examine whether we could replicate previous evidence that humans show a self-bias when learning to get rewards for themselves compared to others^3, 20^, we used data from the placebo level (collapsing across trial blocks). We found that our participants were, on average, more likely to select the option with the higher probability of being rewarded when they were playing for themselves than when they were playing for others (χ^2^(1)=19.459, p_boot_<0.001) (Supplementary Figure S3). Therefore, our findings support the idea that humans show bias towards self-oriented learning as opposed to prosocial learning.

We then proceeded by investigating whether varying doses of intranasal oxytocin impacted on the probability of selecting the higher reward option during self-oriented and prosocial learning. We used a generalized logistic mixed model where we predicted trial-by-trial choices (0 = lowest chance of selecting the reward option; 1 = highest chance of selecting the reward option) using trial number (1-16 within each block), block (1-4), learning condition (self-oriented or prosocial) and treatment level (placebo, low, medium or high) plus all possible interactions as fixed effects and individuals as a random effect.

We found a significant main effect of trial number in predicting trial-by-trial performance (χ^2^ (15) = 733.648, p_boot_ < 0.001; β = 0.079) which suggested that participants, irrespective of learning condition, block or treatment level improved their performance over trials (none of all possible interactions between trial and block, learning condition or treatment were significant; therefore, we excluded all interactions with trial from the final analysis to obtain a more parsimonious model - ΔBIC_full-reduced_ > 100) (Table 1). This analysis further confirmed that participants were able to complete the task successfully.

**Table 1.** Dose-response effects of intranasal oxytocin on self-oriented and prosocial reinforcement learning (generalized logistic mixed model). To investigate dose-response effects of intranasal oxytocin on self-oriented and prosocial reinforcement learning, we used a generalized logistic mixed model where we tried to predict trial-by-trial choices (0 – lower chance of reward option; 1 – higher chance of reward option) using trial number, block, learning condition, treatment and all possible interactions as fixed predictors and participants as random effects. We present a summary of the type III likelihood ratio tests for fixed effects. Significance was assessed with bootstrapping (1000 samples).

| Type III fixed effects |  |  |  |
| --- | --- | --- | --- |
| Effect | df | $\chi^2$ | p (bootstrap) |
| Trial | 15 | 733.648 | < 0.001 |
| Block | 3 | 53.502 | < 0.001 |
| Learning condition | 1 | 138.240 | < 0.001 |
| Treatment | 3 | 6.331 | 0.097 |
| Block * Learning condition | 3 | 151.056 | < 0.001 |
| Block * Treatment | 9 | 21.695 | 0.010 |
| Learning condition* Treatment | 3 | 6.024 | 0.110 |
| Block * Learning condition* Treatment | 9 | 23.382 | 0.005 |

We also found a significant three-way interaction of learning condition x block x treatment (χ^2^ (9) = 23.382, p_boot_ = 0.005) (Table 1). Plotting the data (Figure 2) suggested that the three-way interaction was driven by the following: while participants learnt to get rewards both for themselves and others, performance in the self-oriented learning condition remained high across blocks and treatment levels, while performance in the prosocial learning condition decreased in the last block for all treatment levels except for the low dose (Figure 2). Therefore, intranasal oxytocin can affect processes that maintain prosocial performance at steady levels throughout the task, and that this effect is specific for the low dose.

**Figure 2.**
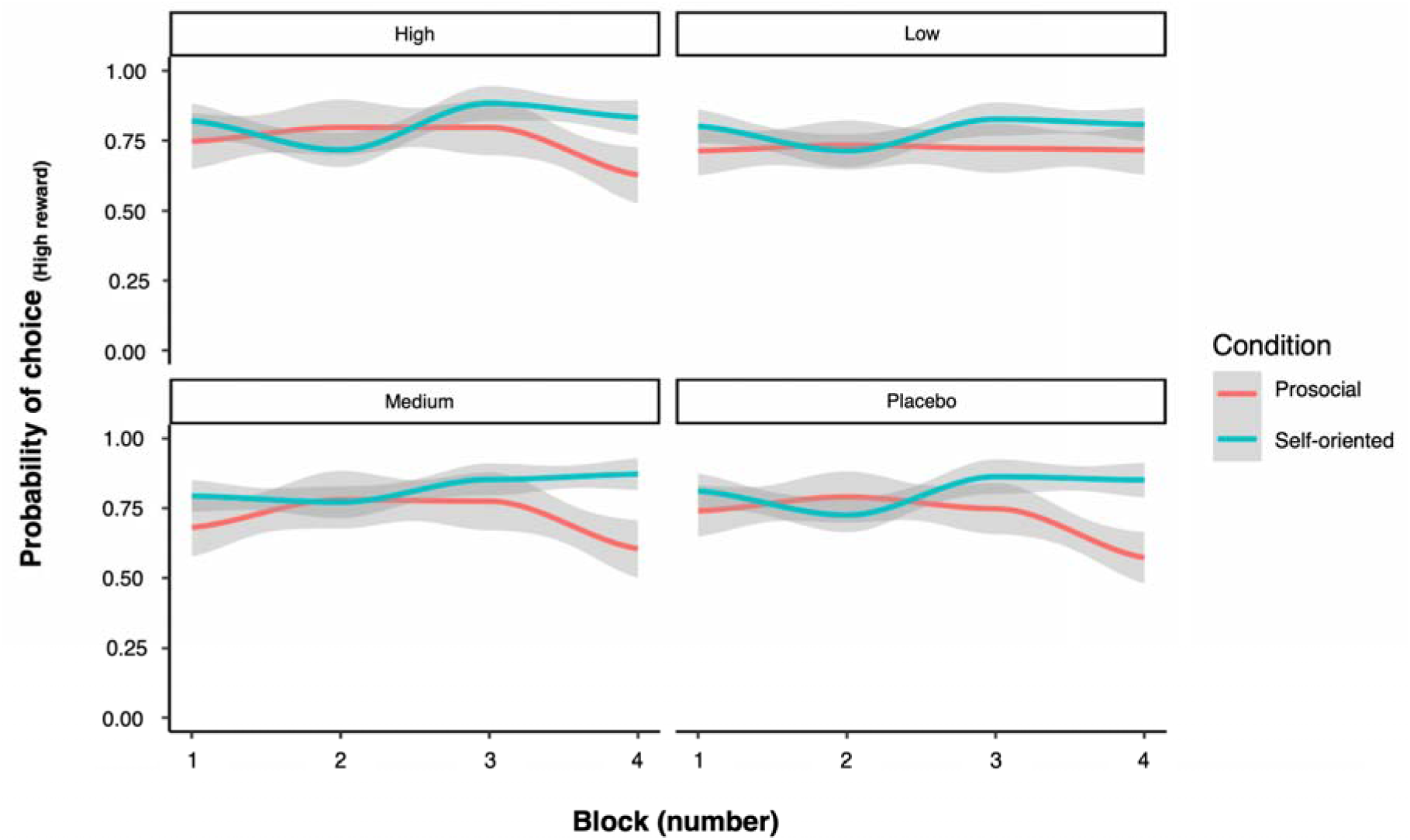
Dose-response effects of intranasal oxytocin on the dynamics of self-oriented and prosocial reinforcement learning over blocks. Evolution across blocks of the probabilities of selecting the option with higher probability of being rewarded, for each treatment and learning condition separately (probabilities were averaged across trials within the same block). Lines result from locally weighted scatterplot smoothing and shades correspond to the respective 95% confidence intervals.

### Behaviour is best explained by a model with separate learning rates for self-oriented and prosocial learning

Next, we used computational models of reinforcement learning to measure two key learning parameters. The learning rate (α) represents the speed at which people update future outcome expectations based on past outcomes. The temperature parameter (β) represents the exploitation - exploration trade-off during action selection, i.e. extent to which the subject decides to stay with what they expect to be the most rewarding option vs exploring other potentially rewarding actions. We modelled learning during the task by fitting five models based on the *Rescorla-Wagner* reinforcement learning algorithm^21^ to data pooled across all treatment levels. The models varied in their combination of α and β parameters they included for each learning condition (Table 2).

**Table 2.**
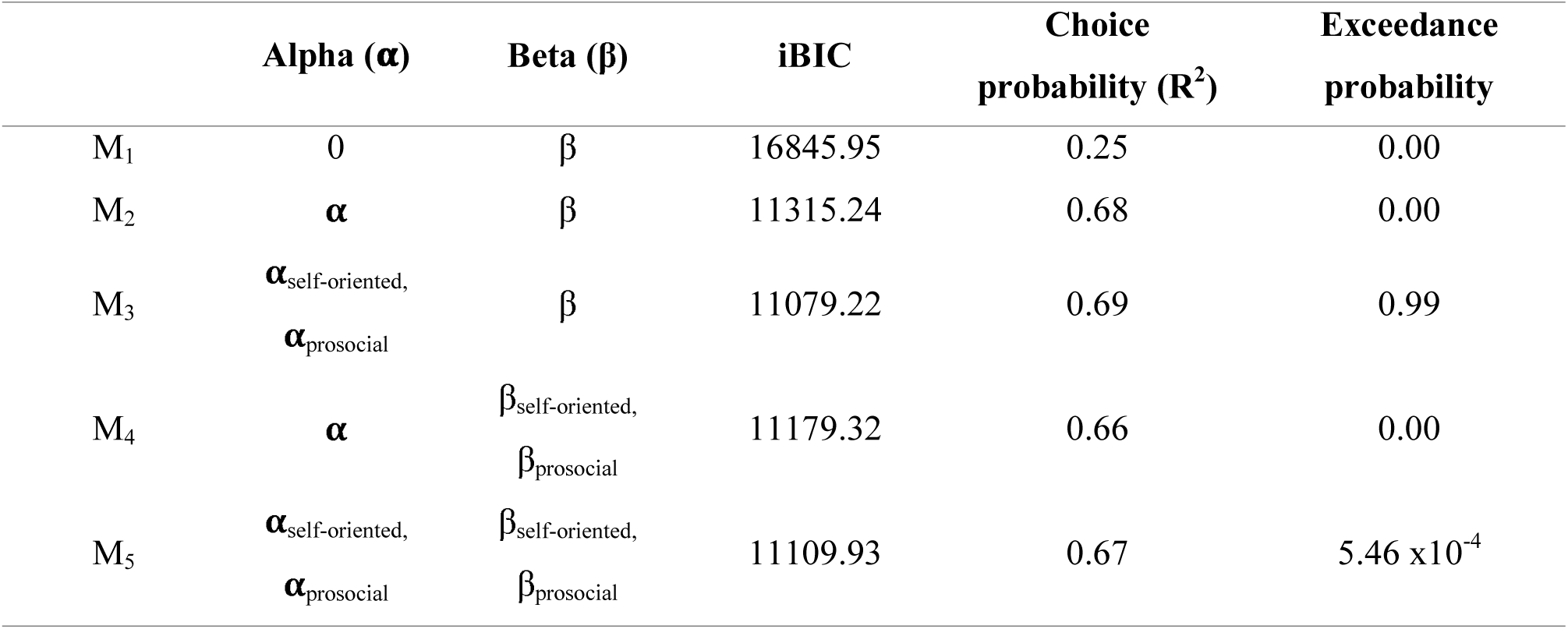
Computational modelling - Model space and selection: We used *Rescorla-Wagner* (RW) computational models of reinforcement learning to estimate learning rates (α) and temperature parameters (β). Our model space included five competing models. In each model, we created variations of the classical RW through the number of parameters used to explain the learning rate and temperature parameters in the task (M_1_-M_5_). We fitted all models pooling data across treatment levels. Our model selection procedure was based on three criteria. First, we used the integrated Bayesian Information Criteria (iBIC) to perform fixed effects model selection (lower is better). Second, we examined the predictive capability of each model in predicting choice probability (higher is better) (R^2^). Third, we performed Bayesian model selection and calculated the exceedance probability of each model (higher is better). M_3_ was the winning model according to the three criteria.

Model selection using both fixed and random effects approaches showed the best model was M_3_, which included different learning rate parameters for self-oriented and prosocial learning (α_self_ and α_prosocial,_ respectively_)_, and one single temperature parameter for both conditions. This model had the lowest integrated BIC (11079.22) highest exceedance probability (0.99) and explained the greatest variance of individual behaviour, among all participants and treatment levels (r^2^ = 0.69; Table 2). M_1_, a null model where we fixed a single α = 0 across learning conditions showed the worst performance, as compared to all the other models where we fitted a learning rate parameter for at least one learning condition (Table 2). This finding strengthens our conclusion that participants successfully learnt the task. We additionally verified the following: i) that parameters in our winning could be estimated independently from each other (Supplementary Figure S4); ii) that our winning model won for each treatment level (Supplementary Figure S5 and Supplementary Table S1); iii) that our winning model was identifiable (Supplementary Figure S6); and iv) that the estimated parameters were recoverable (Supplementary Figure S7). Furthermore, using choices simulated from the *maximum a posteriori* estimates of the parameters previously estimated for each of our participants, we also verified that our winning model could predict their actual choices (r^2^ ranged between 0.224 and 0.841; smallest p = 0.019) (Supplementary Table 2).

### Intranasal oxytocin does not impact the rate at which people learn during self-oriented or prosocial learning

Next, we used the parameters of our validated winning model to test for the effects of learning condition, treatment, and learning condition x treatment on the learning rate and temperature parameters. We found a significant main effect of learning condition (χ^2^ (1) = 5.773, p_boot_ = 0.016), which reflected the fact that our participants showed higher learning rates during self-oriented as compared to prosocial learning (Supplementary Figure S8). This finding is consistent with the self-oriented bias found in performance. The main effect of treatment (χ^2^ (3) = 0.670, p_boot_ = 0.877) and the learning condition x treatment interaction (χ^2^ (3) = 3.016, p_boot_ = 0.403) were not significant. We also tested the main effect of treatment on the temperature parameter, which was not significant (χ^2^ (3) = 2.650, p_boot_ = 0.449) (Supplementary Figure S9).

### Intranasal oxytocin modulates the encoding of prediction errors in the midbrain and sgACC during prosocial learning in a dose-dependent manner

In the RL framework, two quantities are computed during learning: i) expected value of the chosen action at the cue phase (when participants see the options they can choose from); ii) PEs at the feedback phase (when participants receive feedback about whether their choice was rewarded or not). Hence, we investigated whether intranasal oxytocin impacted on brain representations of expected value of chosen actions and PEs during self-oriented and prosocial learning. We used the output of the winning model to estimate these parameters.

First, we used data from the placebo session to examine whether the BOLD signal in three *a priori* defined anatomical regions-of-interest, the nucleus accumbens, the sgACC, and the midbrain, tracked PEs during self-oriented and prosocial learning as hypothesised. We found that PEs for both the self-oriented and prosocial conditions were tracked in the nucleus accumbens (self-oriented condition: mean parameter estimate 0.404 CI95% [0.260, 0.548]; prosocial condition: 0.217 CI95% [0.129, 0.305]) and the midbrain (self-oriented condition: 0.291 CI95% [0.239, 0.343]; prosocial condition: 0.269 CI95% [0.213, 0.325]). The BOLD signal representations of PEs in the nucleus accumbens were stronger in the self-oriented condition than in the prosocial conditions (χ^2^ (1) = 5.033, p_boot_ = 0.026). There was no significant effect of learning condition in the midbrain (χ^2^ (1) = 0.337, p_boot_ = 0.576). Critically, we found that the sgACC specifically encoded PEs in the prosocial but not in the self-oriented conditions (self-oriented condition: -0.056 CI95% [-0.138, 0.026]; prosocial condition: 0.500 CI95% [0.364, 0.636]; self-oriented versus prosocial conditions comparison: (χ^2^ (1) = 34.335, p_boot_ < 0.001) (Supplementary Figure S10). Parameter estimates for the BOLD signal representations of PEs in the prosocial condition in the sgACC correlated positively with inter-individual differences in learning rates in the prosocial condition (r(22) = 0.664, p_boot_ = 0.001), but not in the self-oriented condition (r(22) = 0.349, p_boot_ = 0.103) (Supplementary Figure S11). Direct comparisons of these two correlations yielded no significant differences (Z=-1.378, p=0.084). Parameter estimates for the BOLD signal representations of PEs during the self-oriented and prosocial learning conditions in the nucleus accumbens and the midbrain correlated positively with inter-individual differences in learning rates in both conditions (Nucleus accumbens: self-oriented condition – r(22) = 0.655, p_boot_ = 0.001; prosocial condition – r(22) = 0.590, p_boot_ = 0.003; Self-oriented vs prosocial condition - Z=0.330, p=0.741; Midbrain: self-oriented condition – r(22) = 0.545, p_boot_ = 0.007; prosocial condition – r(22) = 0.594, p_boot_ = 0.003; Self-oriented vs prosocial condition - Z=-0.220, p=0.826;) (Supplementary Figure S11). We also conducted exploratory whole-brain analyses comparing the BOLD signal representations of PEs between the self-oriented and the prosocial learning conditions but no cluster survived correction for multiple comparisons (see Supplementary Figure S12 for brain regions where the BOLD signal tracked PEs in each condition separately).

Next, we tested whether oxytocin impacted on BOLD signal representations of PEs in our three ROIs. We found significant interactions between learning condition and treatment for the sgACC (χ^2^(3)=16.431, p_boot_=0.004) and midbrain (χ^2^ (3)=11.058, p_boot_=0.011) (see Supplementary Table S3 for main effects). In the sgACC, this interaction was driven by an inverted-U-like dose-response like pattern, where the low dose increased the BOLD signal representations of PEs in the prosocial condition, but the high dose decreased the BOLD signal representations, as compared to placebo (Figure 3; please see Supplementary Table S4 for post hoc tests). For the midbrain, we noted the same inverted-U-like dose-response pattern we describe for PEs in the prosocial condition in the sgACC (Figure 3; see Supplementary Table S4 for post hoc tests). None of the three doses of intranasal oxytocin affected the BOLD signal representation of PEs in the sgACC and midbrain during the self-oriented condition (Figure 3). For the nucleus accumbens, only the main effect of learning condition was significant (χ^2^ (1) = 18.803, p_boot_ < 0.001): the BOLD signal representations of PEs in this region were stronger in the self-oriented than the prosocial conditions across treatment levels (Figure 3; Supplementary Table S3).

**Figure 3.**
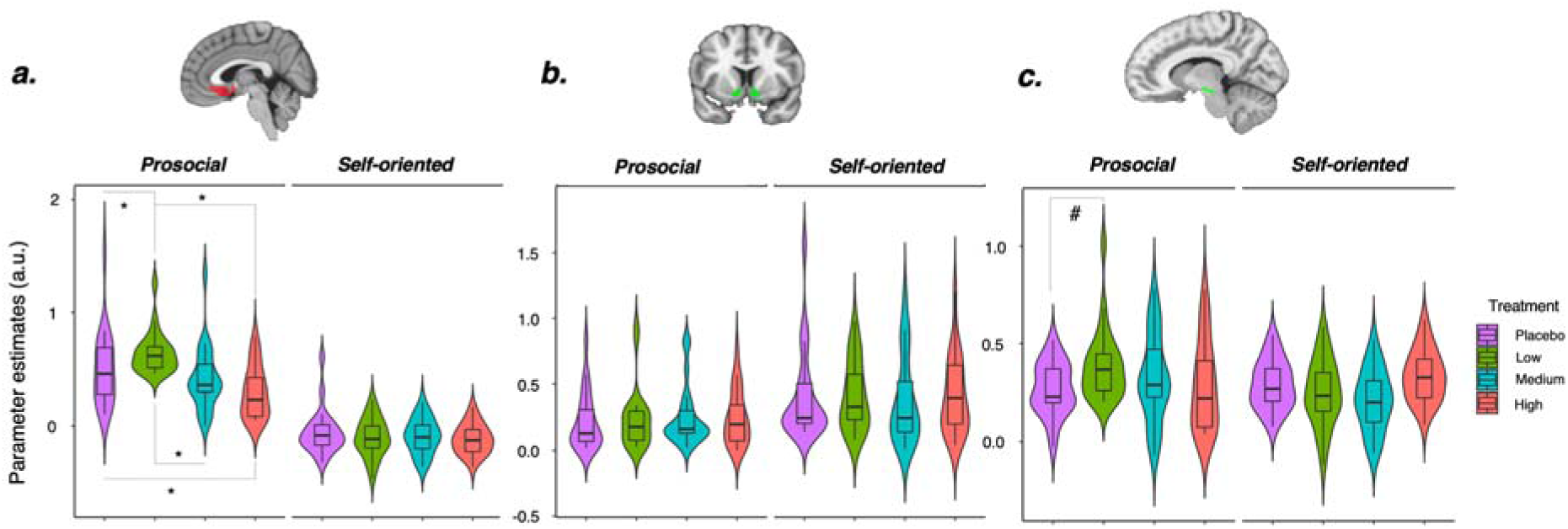
Dose-response effects of intranasal oxytocin on encoding of prediction errors in the subgenual anterior cingulate cortex, nucleus accumbens and midbrain. Learning condition, treatment and learning condition x treatment effects on encoding of prediction errors in the BOLD signal of the subgenual anterior cingulate (a), nucleus accumbens (b) and midbrain (c). Significant interactions were followed up with post hoc tests, applying the Holm-Bonferroni correction for multiple testing. * indicates p_adj_<0.05; # indicates p_adj_ = 0.067 (trend-level).

Since we did not define any strong *a priori* hypothesis about specific brain regions encoding the expected value of the chosen action (at the cue phase), we conducted exploratory whole-brain analyses. We found that the expected value of both self-oriented and prosocial chosen actions was tracked positively by the BOLD signal in a network of areas encompassing the basal ganglia, frontal and occipital cortices and the cerebellum (Supplementary Figure S13). Direct comparisons between the self-oriented and prosocial conditions did not yield significant differences (no cluster survived correction). Intranasal oxytocin did not impact on the BOLD representations of expected values of the chosen actions neither during self-oriented or prosocial learning (no cluster depicting treatment or learning condition x treatment effects survived correction).

### Intranasal oxytocin modulates the encoding of prediction errors in the functional coupling between the midbrain and sgACC during prosocial learning in a dose-dependent manner

Prediction errors are typically encoded in dopaminergic midbrain neurons^6^. The sgACC also receives dense dopaminergic innervation from the midbrain^22, 23^. Hence, it is plausible that the strength of the functional coupling between the midbrain and sgACC might track PEs in the prosocial condition. To test this hypothesis, we used our placebo data to conduct psychophysiological interaction (PPI) analyses with the midbrain as the seed region. We found that the BOLD signal tracking PEs in the midbrain was positively coupled with the BOLD signal in the sgACC in the prosocial learning, but not the self-oriented learning conditions (self-oriented learning: 0.069 CI95% [-0.053, 0.191]; prosocial learning: 0.501 CI95% [0.369, 0.633]; self-oriented versus prosocial learning comparison: (χ^2^ (1) = 27.132, p_boot_ < 0.001); Supplementary Figure S14). The magnitude of the coupling between the BOLD signal tracking PEs in the midbrain and the sgACC was positively correlated with inter-individual differences in learning rates in the prosocial condition (r(22) = 0.859, p_boot_ < 0.001), but not in the self-oriented condition (r(22) = 0.308, p_boot_ < 0.153; self-oriented versus prosocial learning comparison: Z=3.071, p=0.001; Supplementary Figure S15). We also found that the BOLD signal tracking PEs in the midbrain was coupled with the BOLD signal tracking PEs in the nucleus accumbens during both self-oriented and prosocial learning (self-oriented learning: 0.534 CI95% [0.474, 0.594]; prosocial learning: 0.304 CI95% [0.218, 0.390]). However, in this encoding was stronger for self-oriented as compared prosocial learning (χ^2^ (1) = 16.955, p_boot_ < 0.001). The strength to which PEs during self-oriented and prosocial learning were encoded in the functional coupling between these two regions correlated positively with learning rates in both the self-oriented (r(22) = 0.721, p_boot_ < 0.0001) and the prosocial learning conditions (r(22) = 0.682, p_boot_ < 0.001) (Supplementary Figure S15).

We then tested whether these effects were influenced by oxytocin administration (Supplementary Table S5). In the prosocial learning condition, a low dose compared to placebo strengthened the PE-tracking functional coupling between the midbrain and sgACC while the high dose had the opposite effect, weakening the PE-tracking functional coupling between these two regions (in the same way that was observed when analysing each region separately). In contrast, intranasal oxytocin did not impact on the PE-tracking functional coupling between the midbrain and sgACC in the self-oriented learning condition (learning condition x treatment interaction χ^2^(3)=15.727, p_boot_ < 0.001, Figure 4; please see Supplementary Table S6 for post hoc tests.

**Figure 4.**
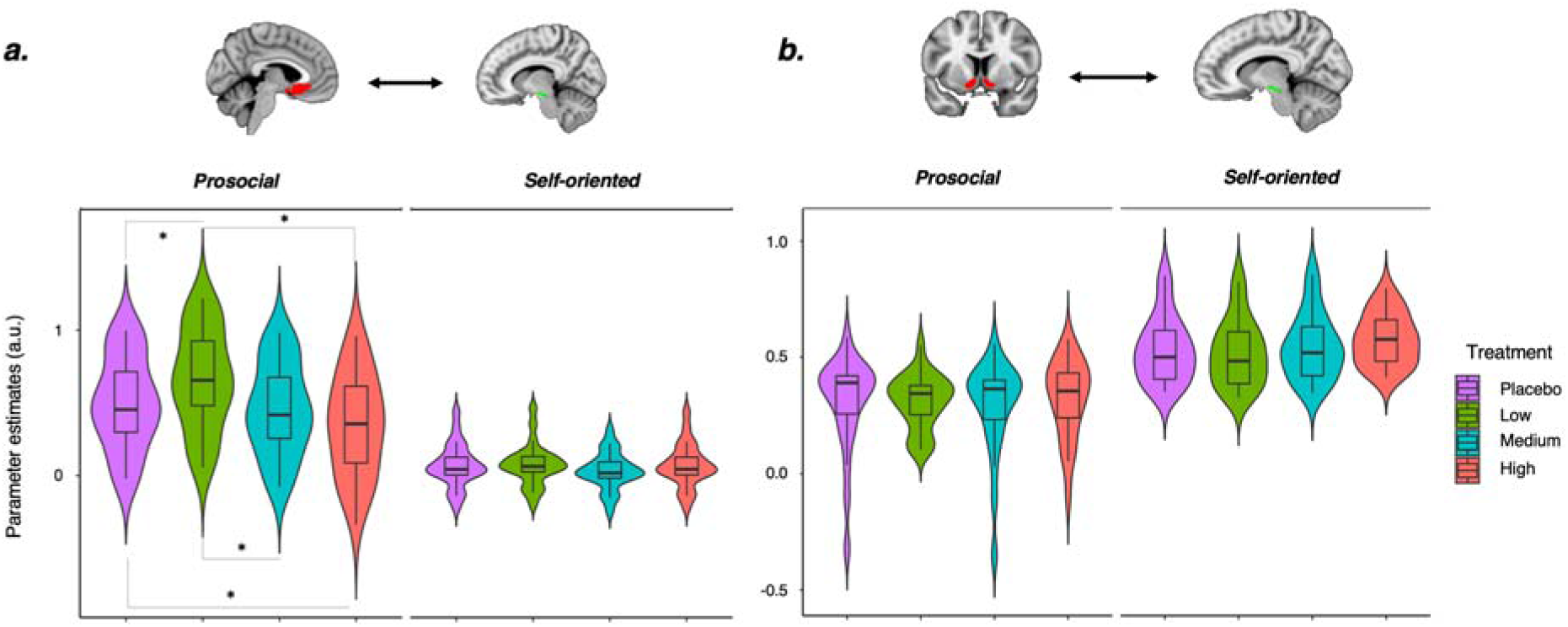
Dose-response effects of intranasal oxytocin on the functional coupling between the midbrain and subgenual anterior cingulate cortex related to prediction errors encoding. Learning condition, treatment and learning condition x treatment effects on psychophysiological interaction parameter estimates reflecting the strength of functional coupling between the subgenual anterior cingulate cortex (a) or the nucleus accumbens (b) and the midbrain associated with encoding of prediction errors during self-oriented and prosocial learning. Significant interactions were followed up with post hoc tests, applying the Holm-Bonferroni correction for multiple testing. * indicates p_adj_<0.05.

For the functional coupling between the midbrain and the nucleus accumbens, only the main effect of learning condition was significant (χ^2^(1)=109.904, p_boot_<0.001). This main effect reflected that the encoding of PEs in the functional coupling between the midbrain and the nucleus accumbens was stronger during self-oriented as compared to prosocial learning (T(22)=12.555, p < 0.001).

### Intranasal oxytocin affects the encoding of PEs in the midbrain-sgACC pathway during prosocial learning by modulating excitatory midbrain-to-sgACC forward transmission and midbrain self-regulation in a dose-dependent manner

Our PPI analyses suggested that intranasal oxytocin modulates the encoding of PE in the midbrain-sgACC pathway during prosocial learning by impacting on the functional coupling strength between these two regions. However, PPI does not provide any information about the direction of this effect^24^. Therefore we conducted dynamic causal modelling (DCM)^25^ to address two questions. First, does intranasal oxytocin modulate the transmission of PE information from the midbrain to the sgACC, vice-versa, or both? Second, does the high dose of intranasal oxytocin decrease the functional coupling between the midbrain and the sgACC by impacting on the intrinsic activity of the midbrain or sgACC, as a result of auto-regulatory mechanisms? For this analysis we used the BOLD signal time series from the midbrain and sgACC regions during the prosocial learning blocks.

We started by defining a fully connected one-state DCM. This full model included forward and backward connections between the midbrain and the sgACC, as well as intrinsic auto-regulatory connections in each node. We used our parametric prosocial PE regressor as input to both nodes. At the first level, we inverted this model for all participants in the four treatment conditions. Commonalities and treatment effects at the group-level were examined within the Parametrical Empirical Bayes (PEB) framework^26^, exploring across all possible reduced PEB models where each parameter or combinations of parameters were switched off one at a time using Bayesian model reduction. To summarize the parameters across all models, we computed the Bayesian model average, which corresponds to the average of the parameters from the top 256 different models, weighted by the model’s posterior probability.

Across all participants and treatment levels, all of our four connections had strong evidence in favour of being different from 0 (posterior probability (P_p_) > 0.95). Our winning second-level model included effects for both the high and low, but not medium dose (Pp = 0.89) (Figure 5). We found strong evidence for decreased intrinsic connectivity in the midbrain after the high dose, as compared to placebo (expected value -0.034, P_p_ = 0.901). Furthermore, we also found strong evidence for increased excitatory transmission from the midbrain to the sgACC after the low dose, as compared to placebo (expected value 0.152, P_p_ = 0.932). Our findings suggest that intranasal oxytocin targets mainly the excitatory connection from the midbrain to the sgACC, whose strength increased after the low dose, compared to placebo, and the intrinsic connectivity of the midbrain, where the high dose produced decreases, as compared to placebo. Finally, we investigated whether the strength of the DCM model connections that were modulated by intranasal oxytocin were predictive of inter-individual differences in prosocial learning. We hypothesised that the excitatory forward connection from the midbrain to the sgACC, where we found dose-dependent modulation by intranasal oxytocin, might be particularly important in explaining inter-individual differences in prosocial learning. We tested this hypothesis by using the PEB modelling procedure described above, but this time testing for correlations between each of our connectivity parameters and learning rates in the self-oriented and prosocial conditions, using the data from the placebo session. We found strong evidence of a positive correlation between the strength of the excitatory forward midbrain-sgACC connection and learning rates in the prosocial condition, but not in the self-oriented condition (expected value 0.120, P_p_ = 0.995; Supplementary Figure S16). We did not find evidence for positive or negative correlations between the strength of any of the other three connections in our model and learning rates, either in the prosocial or the self-oriented learning conditions.

**Figure 5.**
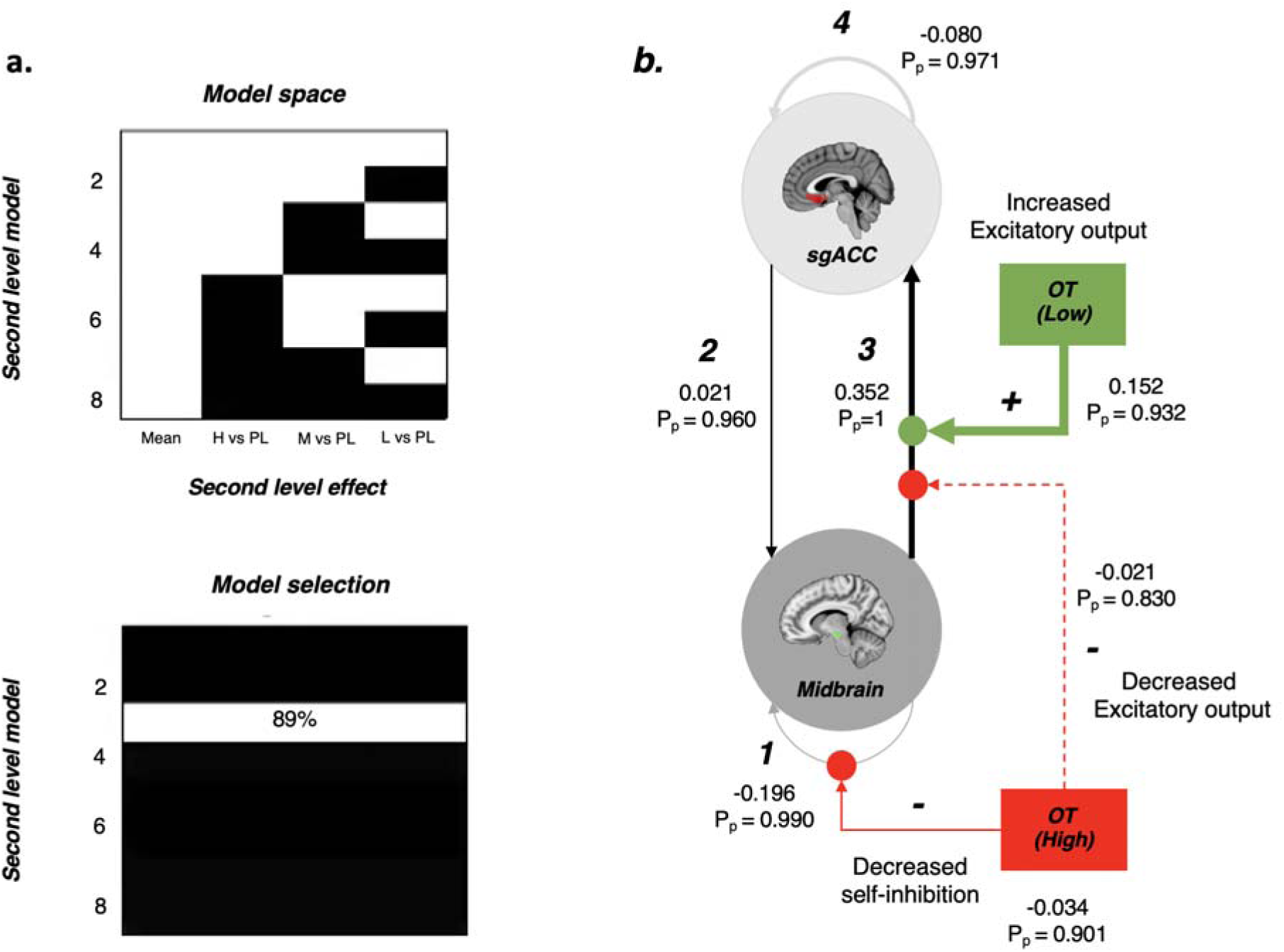
Dose-response effects of intranasal oxytocin on the excitatory midbrain-to-subgenual anterior cingulate (sgACC) forward transmission and midbrain self-inhibition. We conducted dynamic causal modelling (DCM) on BOLD time series from the midbrain and subgenual anterior cingulate cortex (sgACC) during the prosocial blocks to investigate how different doses of intranasal oxytocin modulated effective connectivity between these two regions. We fitted a fully connected one-state vanilla DCM model to all participants and treatment levels at the first-level. We then used the estimates from this first-level models to examine commonalities and treatment effects at the group-level within the Parametric Empirical Bayes framework (second level analysis). Our design matrix for the second level analysis included 4 regressors: i) mean; ii) effects of the low dose as compared to placebo (low vs placebo); iii) medium vs placebo; iv) high vs placebo. Our second level PEB models (a – upper panel) included eight competing models with all possible combinations of treatment effects. M_3_ was the winning model with the highest posterior probability and the lowest free energy (a – lower panel). This model included effects only for the regressors “Low vs placebo” and “High vs placebo”. We investigated these effects further by looking at the expected estimates and posterior probabilities (P_p_) of each parameter of the reduced PEB model. In panel B, we provide a schematic diagram of these effects. Grey/black lines present the mean expected estimates. In green, we present the effects of the low dose; in red, we present the effects of the high dose. Bold lines indicate strong evidence in favour of an expected estimate reliably different from 0 (P_p_>0.90). The dashed line indicates that the evidence was only moderate (P_p_ > 0.80). OT – oxytocin; sgACC – Subgenual anterior cingulate cortex; H – High dose; M – Medium dose; L – Low dose; PL – Placebo; P_p_ – Posterior probability; 1 – Midbrain intrinsic connection; 2 – sgACC – midbrain backwards connection; 3 – Midbrain – sgACC forward connection; 4 – sgAcc intrinsic connection.

## Discussion

We reveal a new role for oxytocin in prosocial learning and its neural mechanisms. First, we demonstrate that a low dose, but not medium or high doses, compared to placebo, can reverse a decrease in performance over time that is specifically observed during prosocial learning (compared to self-oriented learning) during placebo. Second, we demonstrate a dose-dependent modulation in the encoding of PEs in the sgACC, the midbrain, and in the functional coupling between these two regions during prosocial learning, where a low dose strengthens the encoding but a high dose weakens it, compared to placebo. Finally, we demonstrate that the effects of intranasal oxytocin on the encoding of PEs during prosocial learning are likely to emerge from the dose-dependent modulation of both the direct excitatory connections from the midbrain to the sgACC and intrinsic connectivity in the midbrain.

Intranasal oxytocin modulated both performance during learning to benefit others and the neural mechanisms that support prosocial learning but produced no effects on self-oriented learning. Only the low dose of intranasal oxytocin prevented a decrease in learning performance over time that was exhibited in the prosocial (but not the self-oriented) learning conditions under placebo and the two other higher doses. While the exact cognitive mechanisms driving this effect remain elusive, ultimately this effect could result from an amplification of the salience of other-targeted versus self-oriented benefit in response to the administration of a low dose of intranasal oxytocin. This interpretation is supported by previous evidence suggesting that: i) a single dose of intranasal oxytocin increased the willingness to exert effort to get rewards for others in individuals with social anxiety disorder^27^; ii) the effects of intranasal oxytocin on behaviour are likely driven by facilitatory effects on salience processing of social cues^28^.

The lack of an effect on intranasal oxytocin on performance in the self-oriented learning condition contrasts with the findings of a recent behavioural study reporting an overall decrease in self-oriented learning after a single dose of 24 IU of intranasal oxytocin administered with a nasal spray in male and female Chinese students^29^. However, despite the similarity in task design, our studies differ in important methodological aspects, which makes any direct comparison of findings challenging. In addition to differences in the method used for oxytocin administration, there were also marked differences between participant characteristics in the two studies in terms of gender composition and cultural background (our study used only male participants of predominantly white Caucasian ethnicity). Previous evidence has demonstrated that the effects of intranasal oxytocin differ between genders^30^ and cultural backgrounds^31^. To better understand the role that method of administration and participants characteristics may play on the effect of intranasal oxytocin on self-oriented learning behaviour, it is important that future studies systematically investigate these factors.

In addition to uncovering a novel and selective role of oxytocin in prosocial learning, our study provides new insights into differences between learning to benefit the self and learning to benefit others. We showed that performance declined over time in the prosocial learning condition, compared to the self-oriented learning condition. Additionally, we corroborated previous evidence that the same reinforcement learning algorithms provide a foundation both for how humans learn to benefit the self and others^3^. Critically, our findings confirmed that the extent to which these learning mechanisms are invoked is different in self-oriented and prosocial learning, with participants having a higher learning rate for self-oriented reward outcomes compared to reward outcomes benefitting other people^3^.

We provide a detailed map of the brain mechanisms through which intranasal oxytocin modulates prosocial learning by identifying a new and selective role of intranasal oxytocin in the modulation of the encoding of PEs during prosocial learning. Intranasal oxytocin exerted a dose-dependent modulation of the encoding of PEs in the sgACC, the midbrain, and in the functional coupling between these two regions. The selectivity of the effects of intranasal oxytocin in the prosocial condition is congruent with some theories advocating that oxytocin might predominantly affect brain processes related to social functions^28^ (though this idea has been challenged by evidence that intranasal oxytocin also modulates brain functioning during non-social processes^32, 33^). In the context of our task, the selectivity of the effects of intranasal oxytocin in the prosocial condition is intriguing given that both self-oriented and prosocial learning share algorithmic features (both comply with the basic principles of reinforcement learning algorithms and rely on PEs)^5^. Our findings dovetail with a previous study^3^ demonstrating that the way the brain implements reinforcement learning is associated with considerable differences between conditions. For instance, while PEs represent differences between expected and actual outcomes in both self-oriented and prosocial learning, the encoding of PEs during prosocial learning specifically engages the sgACC. Hence, it is plausible that oxytocin might selectively modulate the machinery responsible for the implementation of PE computations during prosocial learning, even if PEs represent the same statistical quantity in both conditions. This idea is further supported by a previous study in rodents showing that the release of oxytocin in the ventral tegmental area increases the excitability of a small subpopulation of neurons engaged during social preference, but not preference for non-social novel objects^13^.

Of particular note is that oxytocin exerted effects on prosocial learning in a manner that was dose-dependent^34^. We found divergent effects for the low and high doses, where a low dose strengthened the encoding of PE (compared to placebo), while a high dose decreased the encoding of PE (compared to placebo). These dose dependent effects of oxytocin are consistent with the effects of intranasal oxytocin on resting regional perfusion in the amygdala in the same cohort of participants^35^. How could different doses of intranasal oxytocin exert opposing effects on the encoding of PEs during prosocial learning in the brain? Oxytocinergic neurons in the hypothalamus project to the midbrain^36, 37^ and facilitate the release of dopamine in the basal ganglia during the encoding of social reward (interacting with a conspecific vs a toy)^13^. Hence, the increase in encoding of PEs during prosocial learning both in the midbrain and sgACC we observed after the low dose could reflect the fact that oxytocin hijacks a population of dopaminergic neurons in the midbrain that project to the sgACC, enhancing the phasic release of dopamine to facilitate the encoding of PEs during prosocial learning^37^. This hypothesis was supported by our DCM analysis, where we found that a low dose of intranasal oxytocin increased the excitatory forward connection from the midbrain to the sgACC - the only connection of our DCM model which explained inter-individual differences in prosocial learning under placebo. At the same time, a high dose of oxytocin might enhance dopamine release to an extent that could result in sustained increases in synaptic dopamine, which in turn might inhibit the release of phasic dopamine through auto-regulatory feedback mechanisms^38^. By inhibiting the release of phasic dopamine, then a high dose of intranasal oxytocin would weaken the encoding of PEs in the prosocial learning condition. In line with this hypothesis our DCM analysis showed reduced intrinsic connectivity in the midbrain after the high dose, as compared to placebo. We note that a similar dose-response model on phasic dopamine release and PE encoding has been shown for amphetamine during self-oriented learning; amphetamine, like oxytocin, also enhances synaptic dopamine^39^.

Our findings also expand our understanding of the neuroanatomical pathways underlying the encoding of PEs during self-oriented and prosocial learning in important ways. First, our results confirm previous evidence suggesting that the sgACC specifically encodes PEs during prosocial learning, while PEs during both self-oriented and prosocial learning are encoded in the nucleus accumbens^3^. However, we expand these findings in two specific ways. We show that PEs during both self-oriented and prosocial learning are similarly encoded in the midbrain. Furthermore, we show that PEs are also encoded in the functional coupling between the midbrain and the sgACC and the nucleus accumbens, in a manner that depends on the recipient of the reward. Prediction errors during prosocial (but not self-oriented) learning are encoded specifically in the functional coupling between the midbrain and sgACC, while PEs during both self-oriented and prosocial learning are encoded in the functional coupling between the midbrain and the nucleus accumbens. Interestingly, we found that the functional coupling between the midbrain and the nucleus accumbens during self-oriented learning is stronger compared to the encoding of PEs during prosocial learning. Hence, the encoding of PEs in the nucleus accumbens exhibited the same self-bias that we observed when we examined performance based on behavioural data and might provide a parsimonious mechanism through which this self-bias emerges.

Even though the BOLD signal is not sensitive to a specific neurotransmitter system, these new findings are consistent with the idea that PEs are encoded in phasic patterns of activity in the dopaminergic neurons of the midbrain, which in turn lead to transient increases in dopamine release in the forebrain, driving synaptic plasticity during learning. The encoding of PEs through phasic dopaminergic activity in the midbrain has mostly been described during self-oriented learning^6^. However, a recent study in rodents showed that dopaminergic neurons in the ventral tegmental area also signal social PEs and drive social reinforcement learning using a similar mechanism^40^. We note that the sgACC also receives dense dopaminergic innervation and expresses receptors for dopamine^23^. Therefore, it plausible that the encoding of prosocial PEs in the midbrain-sgACC pathway might reflect transient patterns of phasic dopamine release from the midbrain into the sgACC and nucleus accumbens, and modulating synaptic plasticity in these regions might produce changes in the BOLD signal. Further studies could test this hypothesis by directly probing phasic dopamine release.

Our study has some limitations that should be acknowledged. First, given the known sexual dimorphism in the oxytocin system, our findings should not be readily extrapolated to women^41–43^. Second, while our findings suggest that oxytocin might interact with the dopamine system to modulate the encoding of PEs, we did not pharmacologically manipulate the dopamine system in this study. This hypothesis is well informed by the known involvement of the midbrain dopaminergic neurons in encoding social PEs^40^ and the engagement of midbrain dopaminergic neurons by oxytocin to encode social reward^13^, but will require further validation in human studies manipulating both systems at the same time. Third, while our dose-response model of the effects of intranasal oxytocin on the encoding of PEs in the prosocial condition suggests that the effects of intranasal oxytocin on the phasic and tonic dopamine release from midbrain neurons to the sgACC may vary by dose, BOLD fMRI does not allow to test this hypothesis directly. This hypothesis could be examined in studies measuring how different doses of oxytocin affect the phasic and tonic dopamine release in the brain during social instrumental learning using voltammetry^44^.

In summary, we demonstrate a new and selective role of intranasal oxytocin in prosocial learning through the modulation of the encoding of PEs in the midbrain-sgACC pathway. Our findings expand our understanding of the neurobiological mechanisms underlying prosocial learning and suggest that dysfunctions in the oxytocin system might play a key role in pathological social behaviour, such as antisocial behaviour, by impeding associative learning of prosocial actions that benefit other people. If that is the case, then oxytocin augmentation might provide an innovative treatment for antisocial behaviour, where we currently lack viable therapeutic options.

## Methods

### Participants

We recruited 24 healthy male adult volunteers (mean age 23.8 years, SD = 3.94, range 20-34 years). We screened participants for psychiatric conditions using the MINI International Neuropsychiatric interview^45^. Participants were not taking any prescribed drugs, did not have a history of drug abuse and tested negative on a urine panel screening test for a range of drugs, consumed <28 units of alcohol per week, and smoked <5 cigarettes per day. We instructed participants to abstain from alcohol and heavy exercise for 24 hours and from food or any beverage other than water for at least 2 hours before scanning. Participants gave written informed consent. King’s College London Research Ethics Committee (HR-17/18-6908) approved the study. We determined sample size based on *a priori* statistical power calculations performed using G*Power (version 3.1). We estimated 24 participants to be the minimally required sample size to detect a within-factor medium effect size of f=0.25 with 80% statistical power (α=0.05) in a repeated measures analysis of variance, assuming a correlation between repeated measures of 0.5.

### Study design

We employed a double-blind, placebo-controlled, crossover design. Participants visited our centre for one screening session and four experimental sessions spaced 4.3 days apart on average (SD = 5.5, range: 2-16 days). During the screening visit, we confirmed participants’ eligibility, obtained informed consent, collected sociodemographic data, and measured weight and height. Participants also completed a short battery of self-report questionnaires (which were collected in relation to other tasks and are not reported here). Participants were trained in a mock-scanner during the screening visit to habituate to the scanner environment and minimize its potential distressing impact. Participants were also trained on the correct usage of the PARI SINUS nebulizer, the device that they would use to self-administer oxytocin or placebo in the experimental visits. Participants were randomly allocated to a treatment order using a Latin square design.

### Intranasal oxytocin administration

Participants self-administered one of three nominal doses of oxytocin (Syntocinon; 40IU/ml; Novartis, Basel, Switzerland). We have previously shown that 40IU delivered with the PARI SINUS nebulizer induce robust regional cerebral blood flow (rCBF) changes in the human brain as early as 15-32 mins post-dosing using a within-subject design^46^. In this study, we decided to investigate dose-response using a range of doses smaller than the 40IU we have previously studied, including a low dose (9IU), a medium dose (18IU) and a high dose (36 IU). Placebo contained the same excipients as Syntocinon, except for oxytocin. Immediately before each experimental session started, a researcher not involved in data collection loaded the SINUS nebulizer with 2 ml of a solution (1 ml of which was self-administered) containing oxytocin in the following concentrations 40 IU/ml, 20 IU/ml and 10 IU/ml or placebo (achieved by a simple 2x or 4x dilution with placebo).

Participants self-administered each dose of intranasal oxytocin or placebo, by operating the SINUS nebulizer for three minutes in each nostril (6 min in total), based on a rate of administration of 0.15-0.17 ml per minute. In pilot work using nebulization on a filter, we estimated the actual nominally delivered dose for our protocol to be 9.0IU (CI 95% 8.67 – 9.40) for the low dose, 18.1IU (CI 95% 17.34 – 18.79) for the medium dose and 36.1IU (CI 95% 34.69 – 37.58) for the high dose. The correct application of the device was validated by confirming gravimetrically the administered volume. Participants were instructed to breathe using only their mouth and to keep a constant breathing rate with their soft palate closed, to minimize delivery to the lungs. The *PARI SINUS* nebuliser (PARI GmbH, Starnberg, Germany) is designed to deliver aerosolised drugs to the sinus cavities by ventilating the sinuses via pressure fluctuations. The SINUS nebuliser produces an aerosol with 3 µm mass median diameter which is superimposed with a 44 Hz pulsation frequency. Hence, droplet diameter is roughly one tenth of a nasal spray and its mass is only a thousandth. The efficacy of this system was first shown in a scintigraphy study^47^. Since the entrance of the sinuses is located near the olfactory region, improved delivery to the olfactory region is expected compared to nasal sprays. One study has shown up to 9.0% (±1.9%) of the total administered dose with the SINUS nebuliser to be delivered to the olfactory region, 15.7% (±2.4%) to the upper nose; for standard nasal sprays, less than 4.6% reached the olfactory region^48^. Participants could not guess treatment allocation above chance (reported in our previous manuscript^35^)

### Procedure

All participants were tested at approximately the same time in the afternoon (3-5 pm) for all oxytocin and placebo treatments, to minimise potential circadian variability in resting brain activity^49^ or oxytocin levels^50^. Each experimental session began with an assessment of vitals (blood pressure and heart rate) and the collection of two 4 ml blood samples for plasma isolation (data not reported here). In the first experimental session, participants were also introduced to a confederate as part of the setup of the prosocial learning task (see below for more details). Then we proceeded with the treatment administration protocol that lasted about 6 minutes in total (Fig. 1). Immediately before and after treatment administration, participants completed a set of visual analog scales (VAS) to assess subjective drug effects (alertness, mood and anxiety) (these data have been reported elsewhere^35^). After drug administration, participants were guided to an MRI scanner, where we acquired a BOLD-fMRI scan during a breath-hold task (lasting 5 minutes 16 seconds), followed by 3 pulsed continuous arterial spin labelling (ASL) scans (each lasting 5 minutes and 22 seconds) (data reported elsewhere^35^), the BOLD-fMRI scan during a prosocial reinforcement learning task (21 minutes) reported here, and a resting-state BOLD-fMRI scan (data not reported yet). We decided to collect the data from the prosocial reinforcement learning task at about 34 – 55 mins post-dosing because we have previously demonstrated robust modulation of rCBF in the basal ganglia (a set of regions engaged during reinforcement learning^9^) after a single dose of 40 IU of intranasal oxytocin administered with the PARI SINUS nebuliser during the same time-interval^46^. When the participants left the MRI scanner, we assessed subjective drug effects using the same set of VAS.

### Prosocial reinforcement learning task

The prosocial learning task is a probabilistic reinforcement learning task designed to separately assess self-oriented (rewards for self) and prosocial learning (rewards for another person) ^3, 20^. On each trial participants had to choose between one of two abstract symbols. One symbol was associated with a high probability (75%) and one was associated with a low probability (25%) of a reward. These contingencies were not instructed so had to be learned through trial and error. The two symbols were randomly assigned to the left or right side of the screen and choices were implemented by pressing one of two buttons that corresponded to the selected symbol. Participants selected a symbol and then received feedback on whether the response was correct, so they learned over time which symbol maximised rewards. Trials were presented in blocks, and each block belonged to one of two conditions. In the self-oriented learning condition, earned points translated into increased payment for the participants themselves. These blocks started with “play for you” displayed and had the word “you” at the top of each screen. In the prosocial learning condition, points translated into increased payment for a second participant, who was a confederate that participants met at the start of the first session (see below). Participants were told that they would never meet the other person again, and that the person was not even aware that an additional financial compensation could arise from participants’ performance. The name of the confederate, gender-matched to the participant, was displayed on these blocks at the start and on each screen (Figure 1). Thus, participants were explicitly aware in each trial who their decisions affected. Stimuli were presented using Presentation (Neurobehavioral Systems – https://www.neurobs.com/).

The experiment was subdivided into eight blocks of 16 trials (4 blocks in each condition). Within each block, participants were presented with 16 pairings of the same two symbols. Symbols were not repeated between blocks or sessions. The 4 blocks in each condition were pseudo-randomly ordered in two playlists, which were randomly allocated to participants in equal proportion. In one of the playlists, participants started by playing a self-oriented learning block, while in the other they started with a prosocial learning block. All participants played the same playlist of the task across the four treatment visits.

Participants received instructions for the prosocial reinforcement learning task and how the points they earned would be converted into money for themselves and for the other participant during the screening session. Instructions included that the two symbols differed in their probability of earning points for participants, but that the side on which they appeared on the screen was irrelevant. Participants then completed one block of practice trials before the main task and were informed that outcomes during the practice block would not affect payment for anyone. We briefly repeated these instructions in the beginning of each experimental visit to confirm that participants still remembered the instructions of the task.

The success of the prosocial reinforcement learning task depends on convincing participants that their performance during the prosocial learning blocks will financially benefit someone else. Therefore, our study included an element of deception, whereby we made participants believe that this other person was a real participant enrolled in a secondary arm of the main study. Unbeknown to the participants, this person was a confederate who did not take part in the study but was part of the research team. We allowed for a short period of interaction between participants and confederates right in the beginning of their first experimental visit to increase the plausibility of our deception. Participants only met the confederate once, in the first session. Interaction between participants and confederates was standardized to make sure all participants had similar experiences. Both participants and confederates were instructed they would be only allowed to greet each other and present their names.

After this short period of interaction, the confederate was guided outside of the room. We then asked participants to fill in an impression scale^19^. This scale measured participants’ perception of the confederate using eight questions assessing similarity, perceived group membership, likeability and attractiveness (see Hein et al.^19^ for further details). For each question, participants were asked to select on a 9-point Likert-scale the number that best represented their thoughts about the confederate (i.e. “How similar to you do you think this person is?”; anchors: 1 - “Extremely”; 9 - “Not at all”). Participants were informed that their responses in this scale would be kept anonymous and that the confederates would also fill the same scale to assess their own impression of the participant.

### Computational modelling

We used a reinforcement learning algorithm to model learning in the task. The basis of the reinforcement learning algorithm is the expectation that each choice a on trial *t* is linked with an expected outcome. The value of the expected outcome on trial t+1, *Q_t_*_+1_(a) is quantified as a function of current expectations *Q_t_(*a) and the prediction error δ*_t_*, which is scaled by the learning rate α:

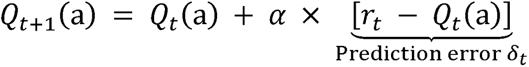

Where δ*_t_*, the prediction error, is the difference between the actual reward experienced on the current trial *r_t_* (1 for reward and 0 for no reward) minus the expected reward on the current trial *Q_t_(*a).

The learning rate α therefore determines the influence of the prediction error. A low learning rate means that new information affects expected value to a lesser extent. The *softmax* link function quantifies the relationship between the expected value of the action *Q_t_(a)* and the probability of choosing that action on trial *t*:

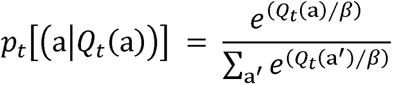

The temperature parameter β represents the noisiness of decisions – whether the participant explores available options or always chooses the option with the highest expected value. A high value for β means that available options are randomly explored as they are equally likely irrespective of their expected value. A low β means that the participant chooses the option with the greatest expected value on all trials. We generated multiple learning models that differed in whether there were separate learning rate and temperature parameters for each learning condition.

### Model fitting

We used MATLAB 2019b (The MathWorks Inc) for all model fitting and comparison. To fit the variations of the learning model (see below) to (real and simulated) participant data we used an iterative maximum a posteriori (MAP) approach as previously described^51, 52^. This method provides a better estimation than a single-step maximum likelihood estimation (MLE) alone by being less susceptible to the influence of outliers. It does this via implementing two levels: the lower level of the individual participants and the higher-level reflecting the full sample. For the real participant data, we fit the model across treatment levels to provide the most conservative comparison, so this full sample combined our four treatment conditions.

For the MAP procedure, we initialized group-level Gaussians as uninformative priors with means of 0.1 (plus some added noise) and variance of 100. During the expectation, we α) for each participant using an MLE approach calculating the log-likelihood of the subject’s series of choices given the model. We then computed the maximum posterior probability estimate, given the observed choices and given the priors computed from the group-level Gaussian, and recomputed the Gaussian distribution over parameters during the maximisation step. We repeated expectation and maximization steps iteratively until convergence of the posterior likelihood summed over the group, or a maximum of 800 steps. Convergence was defined as a change in posterior likelihood <0.001 from one iteration to the next. Bounded free parameters were transformed from the Gaussian space into the native model space via appropriate link functions (e.g. a sigmoid function in the case of the learning rates) to ensure accurate parameter estimation near the bounds.

### Model comparison

We compared five models, which differed in whether the model parameters α and β) for each participant had one value across conditions or varied by the learning condition (self-oriented, prosocial; Table 2). An additional, null model had a learning rate of 0 across both conditions. For model comparison, we calculated the Laplace approximation of the log model evidence (more positive values indicating better model fit) and submitted these to a random-effects analysis using the spm_BMS routine^68^ from SPM12 (http://www.fil.ion.ucl.ac.uk/spm/software/spm12/). This generates the exceedance probability: the posterior probability that each model is the most likely of the model set in the population (higher is better, over 0.95 indicates strong evidence in favour of a model). For the models of real participant data, we also calculated the integrated BIC^51, 52^ (lower is better) and R^2^ as additional measures of model fit. To calculate the model R^2^, we extracted the choice probabilities generated for each participant on each trial from the winning model. We then took the squared median choice probability across participants.

### Simulation experiments

We simulated data from all five models to establish that our model comparison procedure (see above) could accurately identify the best model among the five competing models we included in our model space^20^. For this model identifiability analysis, we simulated 10 datasets including 100 participants, drawing parameters from distributions commonly used in the reinforcement learning literature^53, 54^. Learning rates (α) were drawn from a beta distribution (betapdf(parameter,1.1,1.1)) and *softmax* temperature parameters (β) from a gamma distribution (gampdf(parameter,1.2,5)). We fitted the models to this simulated dataset using the same MAP process as applied to the experimental data from our participants. We then calculated confusion matrices of average exceedance probability (across the 10 runs) and counted how many times each model won.

Our winning model M3 contained three free parameters α_self,_ α_ prosocial,_ β). To assess the reliability of our parameter estimation, we also performed parameter recovery on simulated data as recommended for modelling analyses that use a ‘data first’ approach^55^. We simulated choices 100 times using our experimental schedule and fitted them using the same MAP we described above. We then calculated Pearson’s correlations between the true simulated and fitted parameter values, using bootstrap (1000 samples).

### Statistical analyses of behavioural data

We used one-sample t-tests to investigate: i) whether the mean of the total scores of the impression scale we used to evaluate the perception of the confederates was significantly different from the mid-point of the total score (42); ii) whether participants selected the option with higher probability of being rewarded above chance (0.50) in each condition and treatment level separately. We used a linear mixed-effects model (LMM) to investigate the effect of condition on the probability of selecting the option with higher probability of being rewarded (collapsing across blocks and treatment levels). In this model, we specified condition as a fixed effect and random intercepts for participants. For the trial-by-trial analysis, we used a generalised logistic mixed-effects model to predict binary outcome of choosing the option with the high vs. low probability of being rewarded. The final model did not include any interactions between trial and the remaining factors to obtain a more parsimonious model. For the analysis on learning rates, we used a LMM, where we specified condition, treatment and the interaction between these two factors as fixed effects and random intercepts for participants. For the analysis on the beta parameter, we used a similar LMM, but this time we only specified treatment as a fixed effect. Treatment level was always modelled as a categorical predictor with four levels: placebo, low, medium and high dose. In all models, standard errors and statistical significance were assessed using bootstrapping (1000 samples), as implemented in JASP (version 0.13.1). Significant interactions were followed-up with post-hoc tests, correcting for multiple comparisons with the Holm-Bonferroni procedure.

### MRI data acquisition

We acquired the MRI data in a MR750 3 Tesla GE Discovery Scanner (General Electric, Waukesha, WI, USA) using a 32-channel receive only head coil. We acquired a 3D high-spatial-resolution, Magnetisation Prepared Rapid Acquisition (3D MPRAGE) T1-weighted scan using the following parameters: field of view 270 x 270 mm, matrix = 256 x 256, 1.2 mm thick slices with a slice gap of 1.2 mm, TR/TE/IT = 7312/3016/400 ms, flip angle 11°. The final resolution of the T1-weighted image was 1.1 x 1.1 x 1.2 mm. While participants were performing the prosocial learning task, we acquired functional scans using T2*-sensitive gradient echo planar imaging optimised for parallel imaging, using the following parameters: field of view = 211 x 211 mm, matrix = 64 x 64, 3 mm thick slices with a 3.3 mm slice gap, 41 slices, TR/TE = 2000/30 ms, flip angle = 75°. The final resolution of the functional images was 3.3 x 3.3 x 3.3 mm. The functional imaging sequence was acquired in a descending manner, at an oblique angle (_∼_20°) to the AC–PC line to decrease the effect of susceptibility artifact in the orbitofrontal cortex and midbrain^56^. We also collected field maps (phase-difference B0 estimation; echo time 1 (TE1)□=□4.9□ms, echo time 2 (TE2)□=□7.3□ms) to control for spatial distortions, which are particularly problematic in midbrain fMRI^57^.

### MRI data preprocessing and first-level modelling

#### Preprocessing

We carried out the preprocessing using FEAT, as part of the FMRIB Software Library (FSL) v6.0. Data preprocessing followed a standard pipeline, which included: i) standard head motion correction by volume-realignment to the middle volume using MCFLIRT; ii) distortion correction using phase-difference B0 estimation; iii) slice-time correction; iv) skull-stripping of both functional and structural images using the Brain Extraction Tool (BET); v) high-pass filter (0.01 Hz); vi) registration and spatial normalization to the Montreal Neurological Institute (MNI) 152—*T*_1_ 2-mm template. Individual’s functional images were first registered to their high-resolution MPRAGE scans via a 6-parameter linear registration (FLIRT), and the MPRAGE images were in turn registered to the MNI template via a 12-parameter nonlinear registration (FNIRT). These registrations were combined to align the functional images to the template. Functional images were resampled into the standard space with 2-mm isotropic voxels and were smoothed with a Gaussian kernel of 6-mm full-width at half-maximum. We excluded one participant because they moved excessively in two out of the four sessions (mean frame-wise displacement > 0.5 mm).

#### First-level modelling

We used eight event types to construct regressors in which event timings were convolved with a canonical hemodynamic response function. The two learning conditions at the time of the cues and at the time of the outcome were modelled as separate regressors using stick functions. Each of these regressors was associated with a parametric modulator taken from our winning computational model (M_3_). At the time of the cue this was the value of the chosen action, and at the time of the outcome, the PE. The PEs and values of chosen actions were estimated using mean estimates for alpha and beta across all participants and treatment conditions, calculated for each learning condition separately, as per previous studies^58–60^. This ensures more regularized predictions by minimizing the chance that some participants with smaller alphas will have parametric regressors with very low variance. For all analyses, we mean centred the parametric modulators beforehand and disabled the orthogonalization procedure. This means that all parametric modulators compete for variance, and we thus only report effects that are uniquely attributable to the given regressor. The instruction cue at the beginning of each block was also modelled in a single regressor as a stick function. In some participants, an eighth regressor modelled all missed trials, on which participants did not select one of the two symbols in the response window. We also included 24 head motion parameters (6 head motion parameters, 6 head motion parameters one time point before, and the 12 corresponding squared items) to model the residual effects of head motion as covariates of no interest – this approach has been shown to more efficiently remove head motion effects from BOLD-fMRI data ^61, 62^. We applied pre-withening to remove residual temporal autocorrelation. Subject-level contrast maps were generated using FSL’s FLAME in mixed-effects mode and then used for further second-level analyses, as described below.

### Statistical analyses of fMRI data (second-level)

#### Regions-of-interest analyses

Our ROI analyses were focused on three regions: the sgACC, the nucleus accumbens and the midbrain. In all three regions, we used anatomically defined masks to extract the median parameter estimate of all voxels within each ROI. The sgACC mask included the regions s24 and s25 from the SPM Anatomy toolbox ^63^; the nucleus accumbens and midbrain anatomical masks were derived from a high-resolution atlas of subcortical structures^64^. The midbrain mask included both VTA and SN. These masks were derived from probabilistic anatomical maps by thresholding each map to include voxels with 50% probability or higher of belonging to a certain ROI and then binarizing the thresholded maps. Since the sgACC and nucleus accumbens susceptible to drop out of the BOLD signal^65^, we only extracted data from the voxels of these ROIs that had less than 10% of BOLD signal loss in all participants and treatment conditions. This allowed us to sample within each ROI the same number of voxels in each participants/condition while discarding voxels where the BOLD signal could not be measured reliably. Hence, the final number of voxels in each ROI was: midbrain, 161 voxels; nucleus accumbens = 300 voxels; sgACC = 977 voxels.

We investigated either the effect of learning condition or the effects of learning condition, treatment and learning condition x treatment, as applicable, using LMMs. In all models, we included random intercepts for participants. Significant interactions were followed-up with post-hoc tests, correcting for multiple testing with the Holm-Bonferroni procedure. The correlations between PE parameter estimates in each ROI and learning rates were calculated using Pearson correlation with bootstrapping (1000 samples). Direct comparisons between correlations were performed using the Fisher r-to-Z transform.

#### Whole-brain analyses

We also conducted exploratory analyses at the whole-brain level. For the placebo session where we investigated the effect of learning condition, we performed paired t-tests. For the effects of learning condition, treatment and learning condition x treatment using data from all sessions, we took a partitioned errors approach to account for the likely violation of sphericity present in data from full within-subjects designs^66^. Briefly, to calculate the main effect of learning condition, we averaged the first-level maps across treatment levels for each learning condition and participant and then entered these averaged maps into a paired t-test. To calculate the main effect of treatment, we averaged the first level maps across learning conditions for each treatment level and subject and then entered these averaged maps into a repeated-measures one-way ANOVA. To calculate the learning condition x treatment interaction, we subtracted the first-level maps from learning condition levels and then entered this difference map into a repeated-measures one-way ANOVA. For all whole-brain analyses, we used cluster-level inference at α = 0.05 using family-wise error (FWE) correction for multiple comparisons and a cluster-forming threshold of p=0.001 (uncorrected).

All statistical analyses (behavioural and fMRI data) were conducted with the researcher unblinded regarding treatment condition. Since we used a priori and commonly accepted statistical thresholds and report all observed results at these thresholds, the risk of bias in our analyses is minimal, if not null.

### Psychophysiological interactions (PPI)

We performed psychophysiological interaction analysis^24^ with the midbrain as a seed region. Here, the entire time series over the experiment was extracted from each subject and treatment level from the midbrain anatomical ROI described above. To create the PPI regressors, we multiplied the midbrain time series by the PE parametric regressors. These PPI regressors were used as covariates in a separate PPI-GLM, which included all the regressors plus motion covariates described above for the main first-level GLM. The resulting parameter estimates of the two PPI regressors represent the extent to which activity in each voxel of the brain correlates with the activity in the midbrain that relates to the encoding of PEs during the self-oriented and prosocial learning conditions.

From the individual PPI contrast maps, we extracted the median parameter estimates in all voxels of the sgACC and nucleus accumbens ROIs described above and used these for a number of analyses. First, using data from the placebo session, we tested the effect of learning condition on PE encoding in the functional coupling between the midbrain – sgACC and midbrain – nucleus accumbens. We used LMMs with learning condition as a fixed effect and participant-level random intercepts. Second, we used Pearson correlations with bootstrapping (1000 samples) to investigate correlations between these estimates and the self-oriented and prosocial learning rates. Finally, we investigated learning condition, treatment and learning condition x treatment effects using LMMs, including random intercepts for participants. Significant interactions were followed-up with post-hoc tests, correcting for multiple testing with the Holm-Bonferroni procedure.

### Dynamic Causal Modelling (DCM)

We used a one-state bidirectional DCM model for task fMRI^25^, as implemented in SPM12, to estimate the effective connectivity between the midbrain and sgACC and within each region during the prosocial learning blocks. DCM for fMRI couples a bilinear model of neural dynamics with a biophysical model of hemodynamics to infer effective connectivity between cortical regions^25^. Details regarding this method can be found elsewhere^25^. We extracted the principal eigenvariate of the time-series of the BOLD signal during the prosocial blocks from all voxels in the sgACC and midbrain ROIs, adjusted for the F-contrast of the effects of interest. We defined a fully connected vanilla DCM model, which included both forward and backward connections between the midbrain and sgACC and intrinsic connections within each node. We set PEs as a driving input to both nodes. This full model was inverted for all participants and treatment levels.

The participant-specific DCMs were taken to a second level analysis where we used the Parametrical Empirical Bayes (PEB) approach^26^ as implemented in SPM12 for group level inference; these routines assess how individual (within-subject) connections relate to group means, taking into account both the expected strength of each connection and the associated uncertainty. This means that participants with more uncertain parameter estimates are downweighted, while participants with more precise estimates have greater influence. The PEB approach involves (i) estimating group level parameters using a general linear model (GLM) that divides inter-subject variability into regressor effects and unexplained random effects, followed by (ii) comparison of different combinations of these parameters to identify those that best explain commonalities and differences in connectivity (Bayesian model comparison). Our second level PEB model included four regressors: i) commonalities; ii) effect of low dose (low dose versus placebo); iii) effect of medium dose (medium dose versus placebo); effect of high dose (high dose versus placebo). Each treatment effect regressor specified the placebo condition as -1 and the treatment conditions as 1, so that all regressors were mean centred and the first commonalities regressor estimated the mean group effect. Next, we used Bayesian model reduction (BMR) to test all nested models within each full PEB model (assuming that a different combination of connections could exist for each participant) and to “prune” connection parameters that did not contribute to the model evidence. The parameters of the best 256 pruned models were averaged and weighted by their evidence (Bayesian model averaging, BMA) to generate group estimates of connection parameters. Last, we compared models using free energy and calculated the posterior probability for each model as a *softmax* function of the log Bayes factor. We characterized the between-condition effects on each parameter by using the BMA expected values for the strength of each connection and their respective posterior probability (Pp) of being different from zero. The higher the Pp, the greater the confidence that a certain parameter is different from zero. Here, we interpreted Pp>0.90 as strong evidence and Pp>0.80 as moderate evidence in favour of a reliable difference from zero.

In a secondary analysis, we used data from the placebo session only to investigate whether the strength of the connections in our DCM model could capture inter-individual differences in prosocial learning. We used the DCMs and PEB modelling procedure described above, but this time testing for correlations between each of our connectivity parameters and learning rates during self-oriented and prosocial learning. Hence, our second level PEB GLM model contained three regressors: i) commonalities; ii) mean centred regressor of the learning rates during self-oriented learning; iii) mean centred regressor of the learning rates during prosocial learning.

### Data availability

Data can be accessed from the corresponding author upon reasonable request. The code used for the computational modelling can be found in https://doi.org/10.17605/OSF.IO/XGW7H. A reporting summary for this article is available as a Supplementary Information file.

## List of Supplementary Materials

Supplementary Figure S1. Global impression ratings of the confederates.

Supplementary Figure S2. Task performance during self-oriented and prosocial reinforcement learning.

Supplementary Figure S3. Effect of condition on learning performance during the placebo session.

Supplementary Figure S4. Correlations between the *maximum a posteriori* of the parameters from the winning model M_3_.

Supplementary Figure S5. Bayesian model selection in each treatment level.

Supplementary Table S1. Bayesian model selection in each treatment level (exceedance probabilities).

Supplementary Figure S6. Model identifiability.

Supplementary Figure S7. Parameter recovery of the winning model M_3_.

Supplementary Table S2. Ability of the winning model M_3_ to predict actual behaviour.

Supplementary Figure S8. Effect of condition on learning rates.

Supplementary Figure S9. Effects of treatment on the inverse temperature parameter beta (β).

Supplementary Figure S10. BOLD representations of prediction errors in the subgenual anterior cingulate, nucleus accumbens and midbrain.

Supplementary Figure S11. Correlations between encoding of prediction errors and learning performance.

Supplementary Figure S12. BOLD representations of prediction errors during self-oriented and prosocial learning (whole-brain analysis).

Supplementary Figure S13. BOLD representations of expected value of the chosen actions during self-oriented and prosocial learning (whole-brain analysis).

Supplementary Table S3. Effects of learning condition, treatment and learning condition x treatment on encoding of prediction errors in the subgenual anterior cingulate cortex, nucleus accumbens and midbrain.

Supplementary Table S4. Effect of learning condition x treatment on encoding of prediction errors in the subgenual anterior cingulate cortex and midbrain during self-oriented and prosocial learning (post hoc tests).

Supplementary Figure S14. Encoding of prediction errors in the midbrain functional coupling.

Supplementary Figure S15. Correlations between encoding of prediction errors in the functional coupling of midbrain and performance during self-oriented and prosocial learning.

Supplementary Table S5. Effects of learning condition, treatment and learning condition x treatment on encoding of prediction errors in the functional coupling between the midbrain and the subgenual anterior cingulate cortex, and between the midbrain and nucleus accumbens.

Supplementary Table S6. Effect of learning condition x treatment on encoding of prediction errors in the functional coupling between the midbrain and subgenual anterior cingulate cortex (post hoc tests).

Supplementary Figure S16. Associations between learning rates and the effective connectivity between the midbrain and subgenual anterior cingulate cortex during the prosocial blocks.

## ACKNOWLEDGMENTS

We would like to thank all volunteers contributing data to this study and Uwe Schusching for his help in estimating the actual delivered doses with the nebuliser.

## Funding

This study was part-funded by: an Economic and Social Research Council Grant (ES/K009400/1) to YP; scanning time support by the National Institute for Health Research (NIHR) Biomedical Research Centre at South London and Maudsley NHS Foundation Trust and King’s College London to YP; an unrestricted research grant by PARI GmbH to YP.

## Author contributions

YP, DM and PL designed the study; DM collected the data; DM, PL, JC and RM analyzed the data; DM and YP wrote the first draft of the paper and all co-authors provided critical revisions.

## Competing interests

The authors declare no competing interests. This manuscript represents independent research. The views expressed are those of the authors and not necessarily those of the NHS, the NIHR, the Department of Health and Social Care, or PARI GmbH.

## Notes

### Competing Interest Statement

The authors have declared no competing interest.

