## Supplementary for "Oxytocin modulates neurocomputational mechanisms underlying prosocial reinforcement learning"

**Supplementary Figure S1. Global impression ratings of the confederates.** We assessed participants’ impression of the confederate right after they interacted in the first visit using an impression scale from Hein et al.^1^ and examined whether the confederates might have elicited any form of strong preference bias. In this plot, we show the distribution of the total impression scores of our participants. We tested for statistical differences from the mean total score of the scale (red dashed line) using a one-sample t-test.

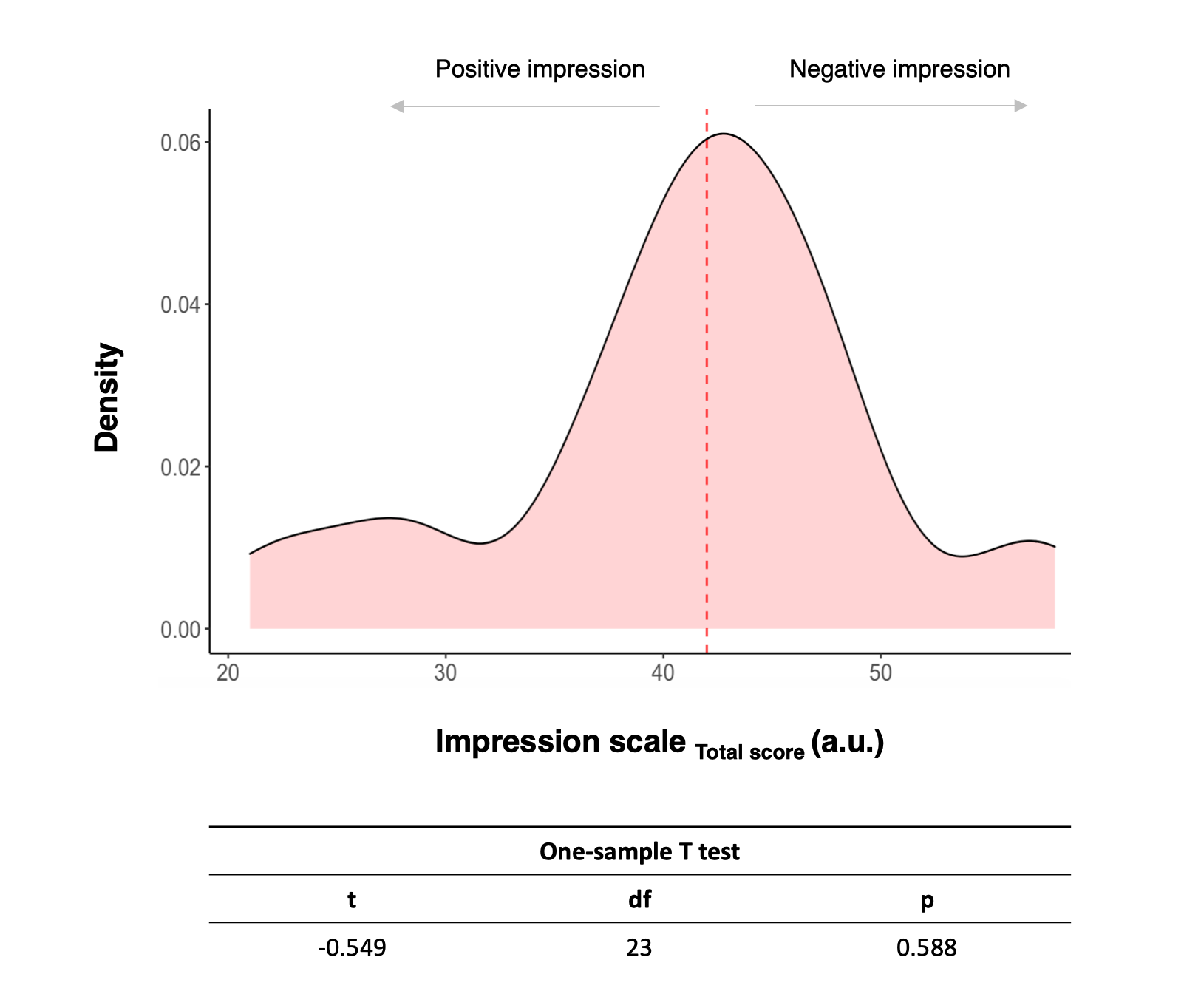

**Supplementary Figure S2. Task performance during self-oriented and prosocial reinforcement learning.** We examined whether participants were able to learn for both recipients (self and other – prosocial learning condition) to validate their ability to complete the task. We quantified performance as selecting the option associated with the higher chance of receiving a reward. (a) We show the results of a set of one-sample t-tests investigating whether participants performed the task above chance (0.5), separately for each learning condition and treatment levels. (b) We provide the corresponding density plots. The dashed line indicates performance at the chance-level (0.5).

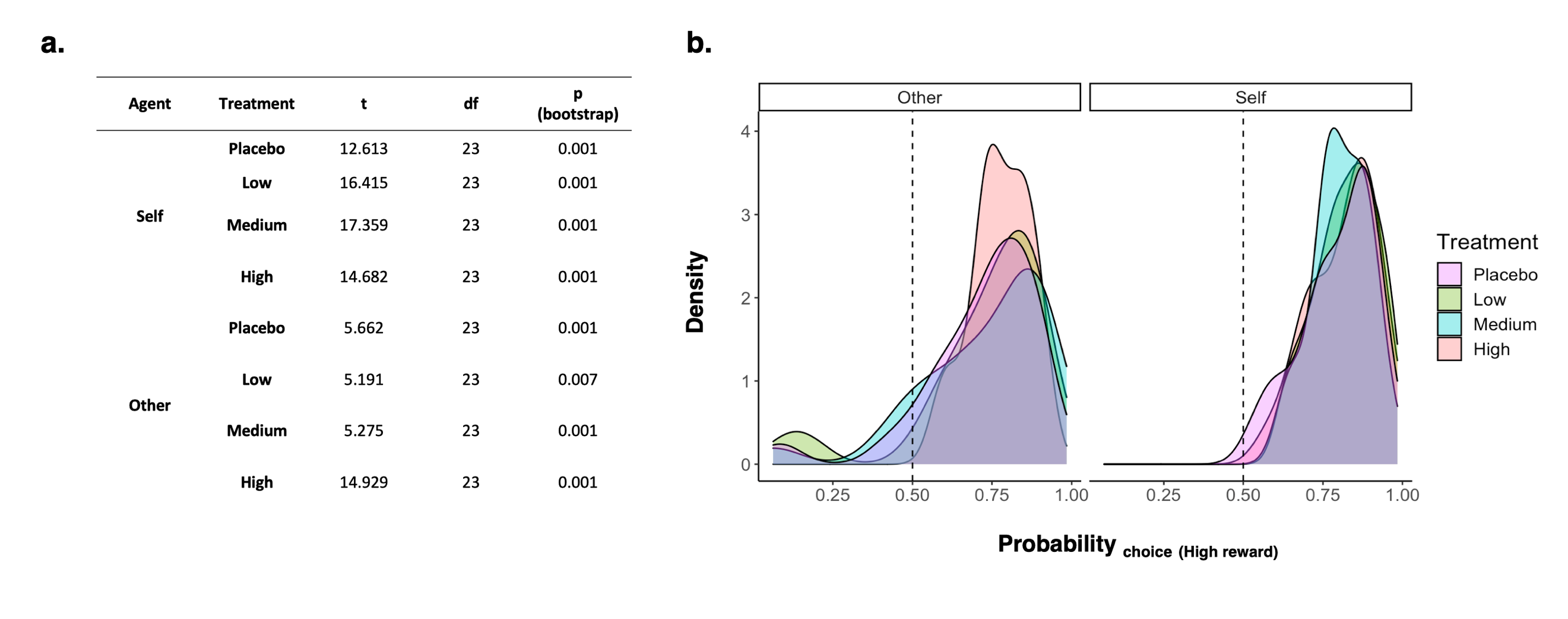

**Supplementary Figure S3. Effect of learning condition on learning performance during the placebo session.** We quantified learning performance using the probability of selecting the option with higher probability of being rewarded averaged across the four blocks of each participant. (a) Learning is shown through evolution across trials of these probabilities for each learning condition separately. Lines depict locally estimated scatterplot smoothing (LOESS) and shadows the respective 95% confidence intervals. (b) Violin plots show performance averaged across trials as well as blocks was higher for self than other. This effect of learning condition was significant in a linear mixed-effects model with random intercepts for participants.

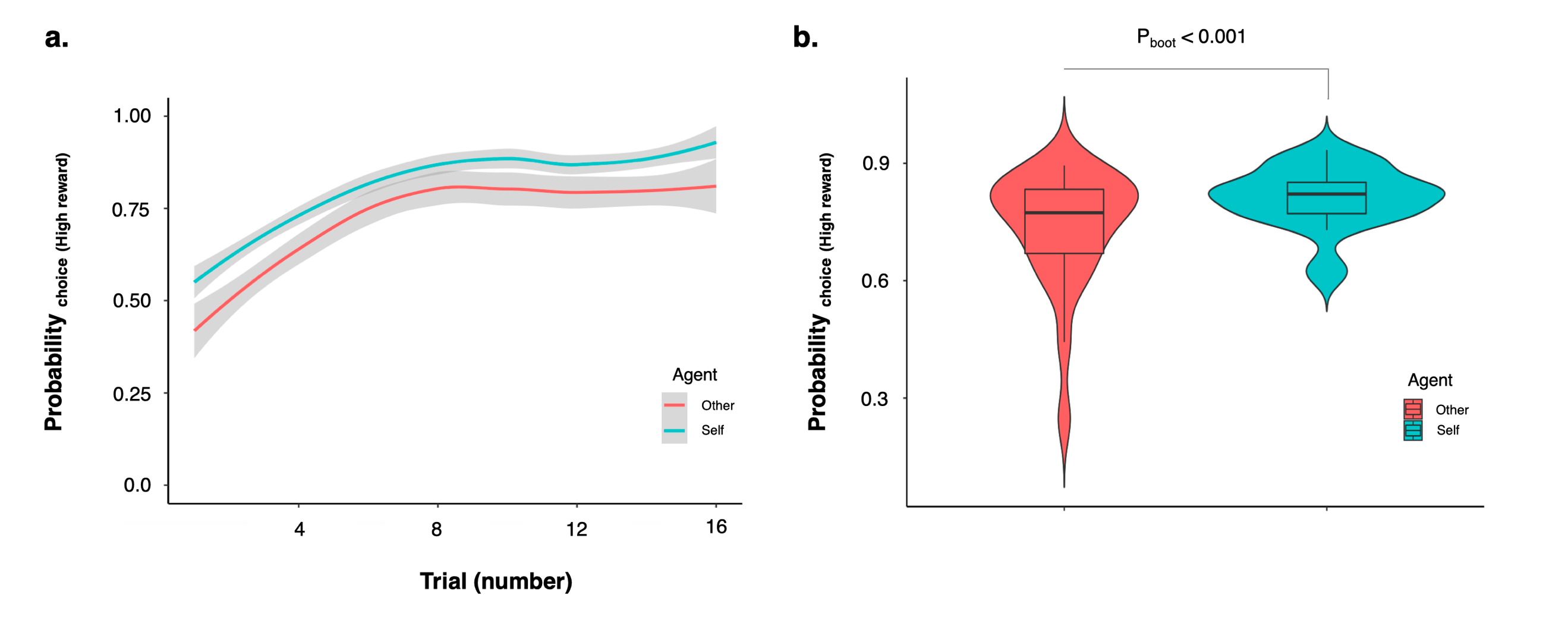

**Supplementary Figure S4. Correlations between the *maximum a posteriori* of the parameters from the winning model M_3_.** To examine whether the *maximum a posteriori (MAP)* of the three parameters in our winning model M_3_ could be estimated independently from each other, we calculated all possible pairwise Pearson’s correlations between our estimates of these parameters across participants. Small correlations indicate that the parameters can be estimate independently from each other. The colour scale depicts the Pearson’s correlation coefficient (r).

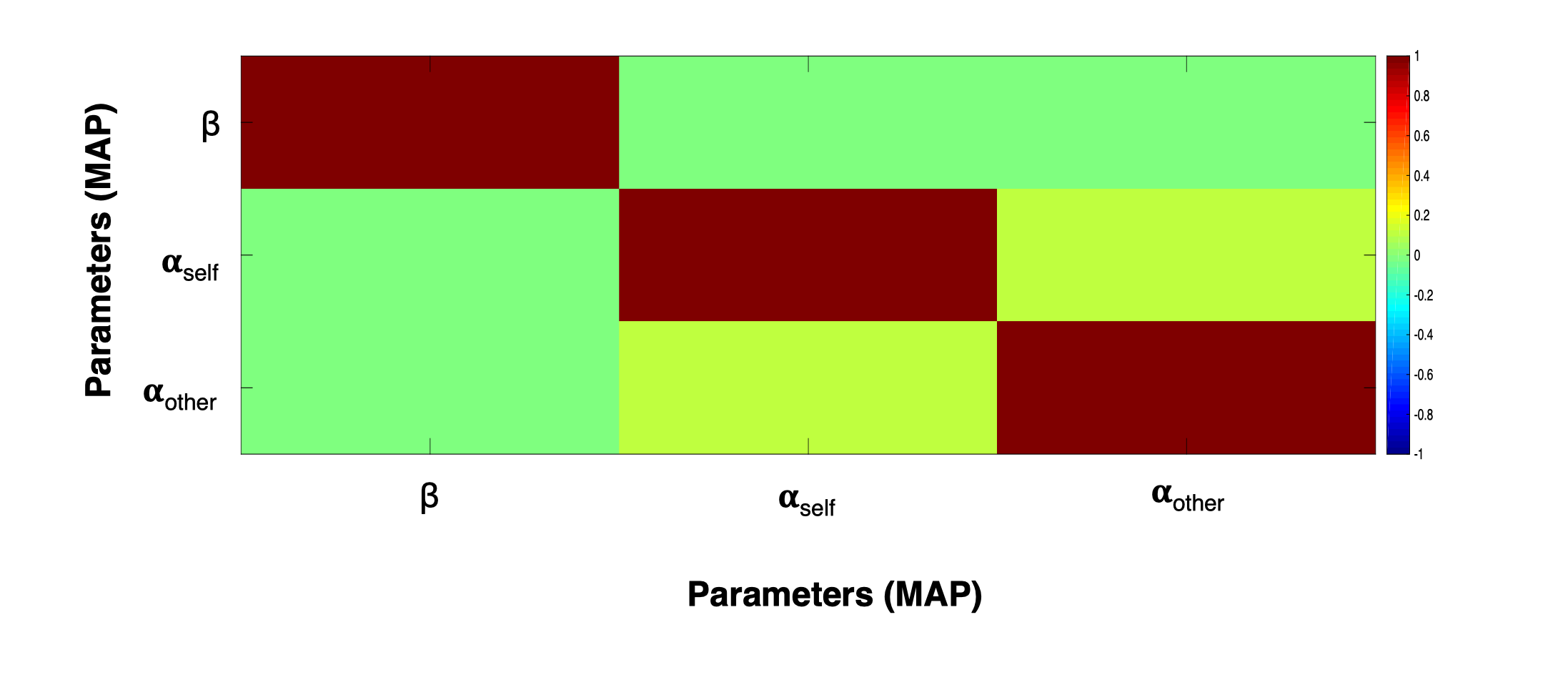

The three parameters could be estimated independently from each other, as suggested by correlations between parameters around 0 (Supplementary Figure S4).

**Supplementary Figure S5. Bayesian model selection in each treatment level.** Here we show the results of the Bayesian model selection performed in each treatment level separately. Using data from each treatment level separately, we calculated the exceedance probability of each of our five models using the same approach we used in our main model selection procedure. M_3_ was consistently selected as the winning model in all treatment levels.

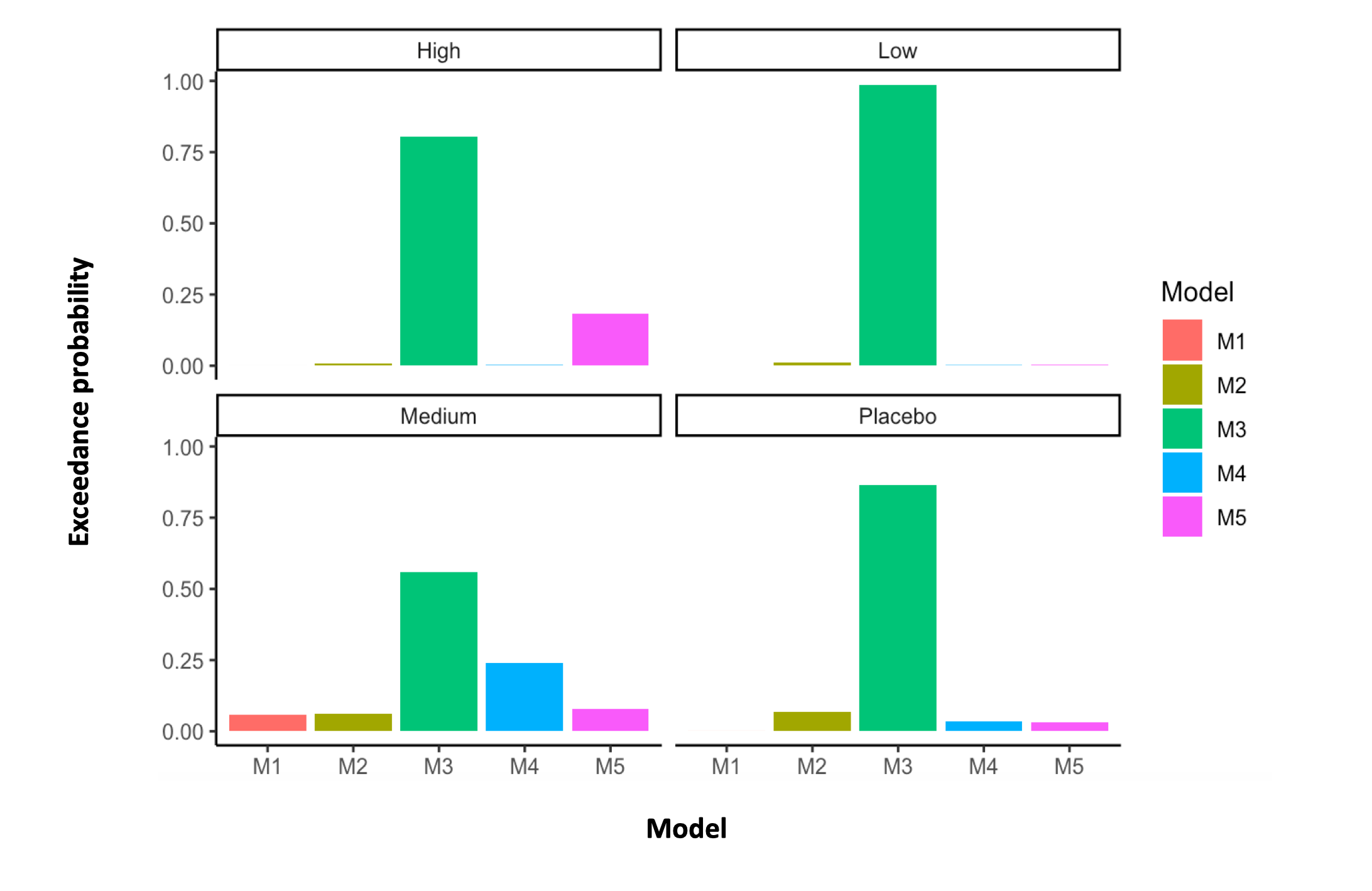

**Supplementary Table S1. Bayesian model selection in each treatment level (exceedance probabilities).** In this table, we provide the exact exceedance probabilities of each model in each treatment level separately.

| **Treatment** | **Model** | **Bayesian Model Selection** |
| --- | --- | --- |
|  |  | **Exceedance probability** |
| **High** | M_1_ | 5.500x10^-5^ |
|  | M_2_ | 0.008 |
|  | M_3_ | 0.805 |
|  | M_4_ | 0.001 |
|  | M_5_ | 0.184 |
| **Medium** | M_1_ | 0.059 |
|  | M_2_ | 0.063 |
|  | M_3_ | 0.558 |
|  | M_4_ | 0.239 |
|  | M_5_ | 0.080 |
| **Low** | M_1_ | 2.300x10^-5^ |
|  | M_2_ | 0.012 |
|  | M_3_ | 0.984 |
|  | M_4_ | 0.001 |
|  | M_5_ | 0.002 |
| **Placebo** | M_1_ | 1.860x10^-4^ |
|  | M_2_ | 0.068 |
|  | M_3_ | 0.864 |
|  | M_4_ | 0.035 |
|  | M_5_ | 0.032 |

We found that M_3_ consistently won in each treatment level (Supplementary Figure S5 and Supplementary Table 1). As a final sanity check, we formally tested whether the same model describes our participants’ behaviour across treatment levels, and we found strong evidence to support this hypothesis (exceedance probability of the null hypothesis of no differences in model frequencies between treatment levels = 1).

**Supplementary Figure S6. Model identifiability.** We created synthetic choices using simulations based on each of our five models, fitted the models to the synthetic data and examined whether the best fitting model was the one that had been used to create the data. This procedure was repeated 10 times. (a) We show the model identifiability average exceedance probability confusion matrix. (b) We show the model identifiability best model selection confusion matrix.

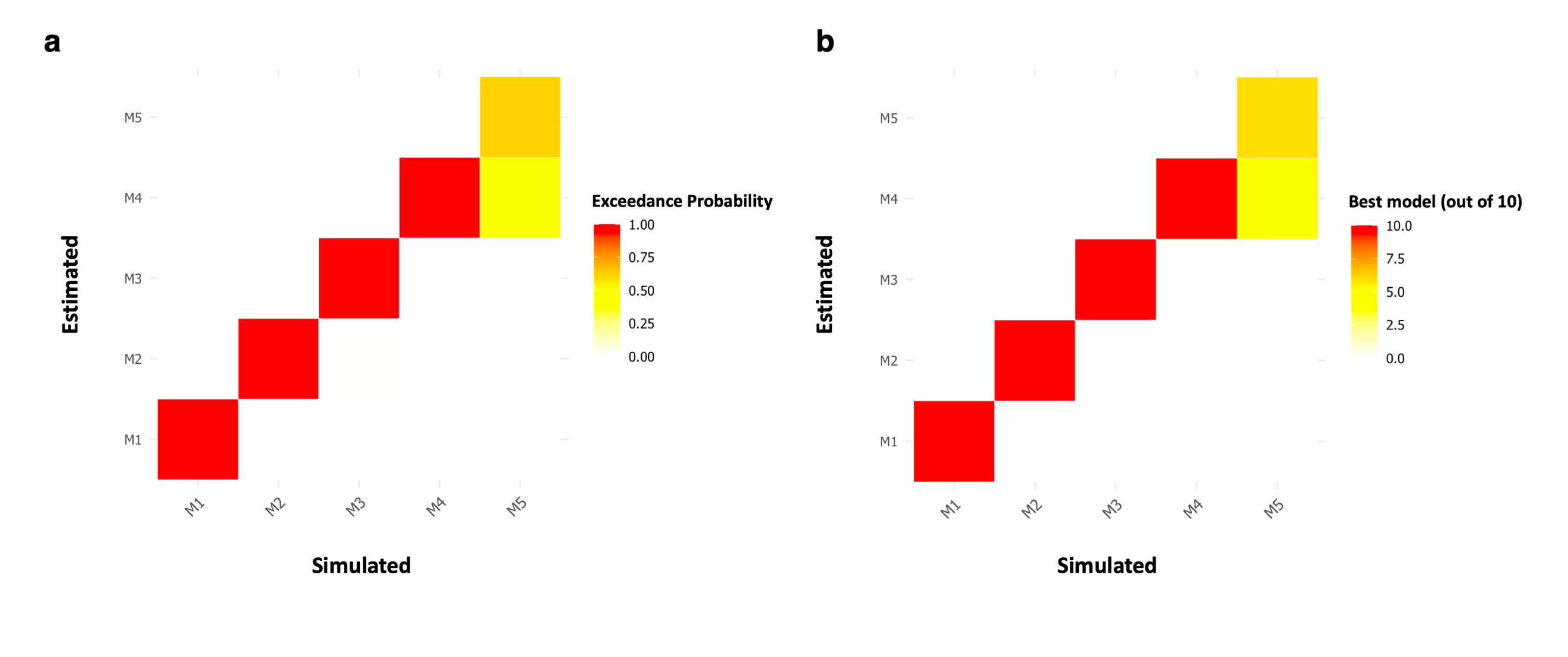

We found that our task design would allow us to clearly identify M_1-4_; M_5_ was still the winning model in most of our simulations, however in 4/10 simulations, M_4_ was selected as winning model even if data was generated from M_5_. Hence, our task design was appropriate to identify our winning model M_3_.

**Supplementary Figure S7. Parameter recovery of the winning model M_3_.** We used our winning model M_3_ to simulate data from 100 participants. Learning rates (α) were drawn from a beta distribution (betapdf(parameter,1.1,1.1)) and softmax temperature parameters (β) from a gamma distribution (gampdf(parameter,1.2,5)) to cover a wide range of parameters estimates. We then fitted M_3_ to the simulated data and recovered the correspondent *maximum a posteriori* (MAP) parameter estimates. To assess parameter recoverability, we calculated Pearson’s correlations (with bootstrap 1000 samples) between the true and recovered parameters. Large correlations indicate good parameter recoverability.

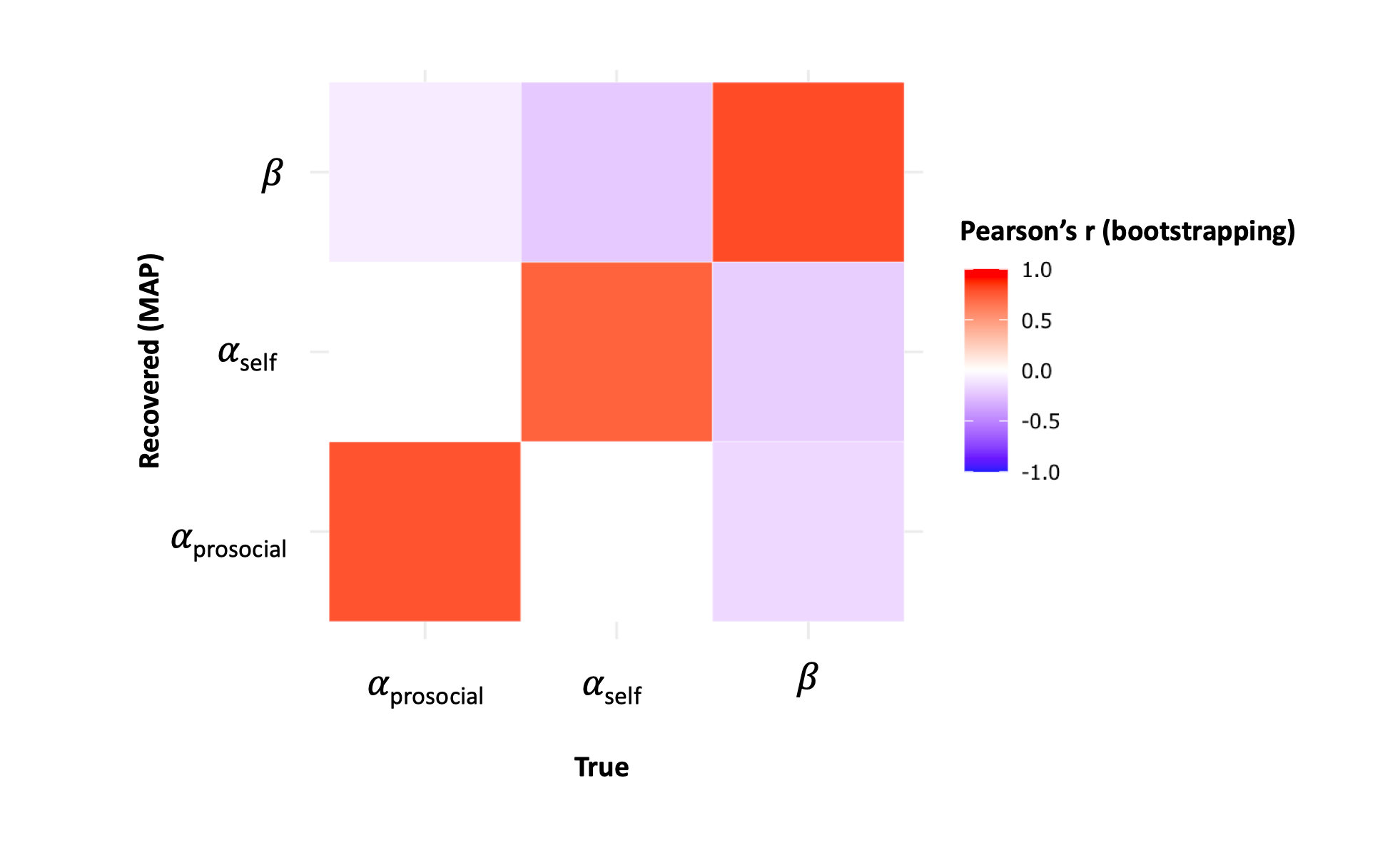

We found high correlations between fitted and recovered parameters from this model (α_self_: r_pearson_ = 0.725 CI95% [0.616, 0.806], p_boot_ < 0.001; α_prosocial_: r_pearson_ = 0.775 CI95% [0.683, 0.843], p_boot_ < 0.001; β: r_pearson_ = 0.797 CI95% [0.713, 0.859], p_boot_ < 0.001).

**Supplementary Table S2. Ability of the winning model M_3_ to predict actual behaviour.** We simulated data from 24 participants using the exact parameter estimates of each of our 24 participants in each of the four treatment levels. From the simulated responses, we calculated the mean probabilities of selecting the option with higher probability of being rewarded and examined whether these simulated probabilities could predict the actual probabilities of our participants in each treatment level. This would provide us with an idea about whether our winning model M_3_ can reliably capture the core features of the actual behaviour of our participants.

| **Treatment** | **Condition** | **r^2^** | **p-value** |
| --- | --- | --- | --- |
| Placebo | Self-oriented | 0.841 | <0.001 |
| Placebo | Prosocial | 0.509 | <0.001 |
| Low | Self-oriented | 0.779 | <0.001 |
| Low | Prosocial | 0.461 | <0.001 |
| Medium | Self-oriented | 0.420 | 0.001 |
| Medium | Prosocial | 0.512 | <0.001 |
| High | Self-oriented | 0.561 | <0.001 |
| High | Prosocial | 0.224 | 0.019 |

**Supplementary Figure S8. Effect of condition on learning rates.** We examined the effects of learning condition, treatment and learning condition x treatment on learning rates, using a linear mixed model where we specified random intercepts for subjects. Only the main effect of learning condition was significant, which we illustrate in this violin plot. Significance was assessed using bootstrap (1000 samples).

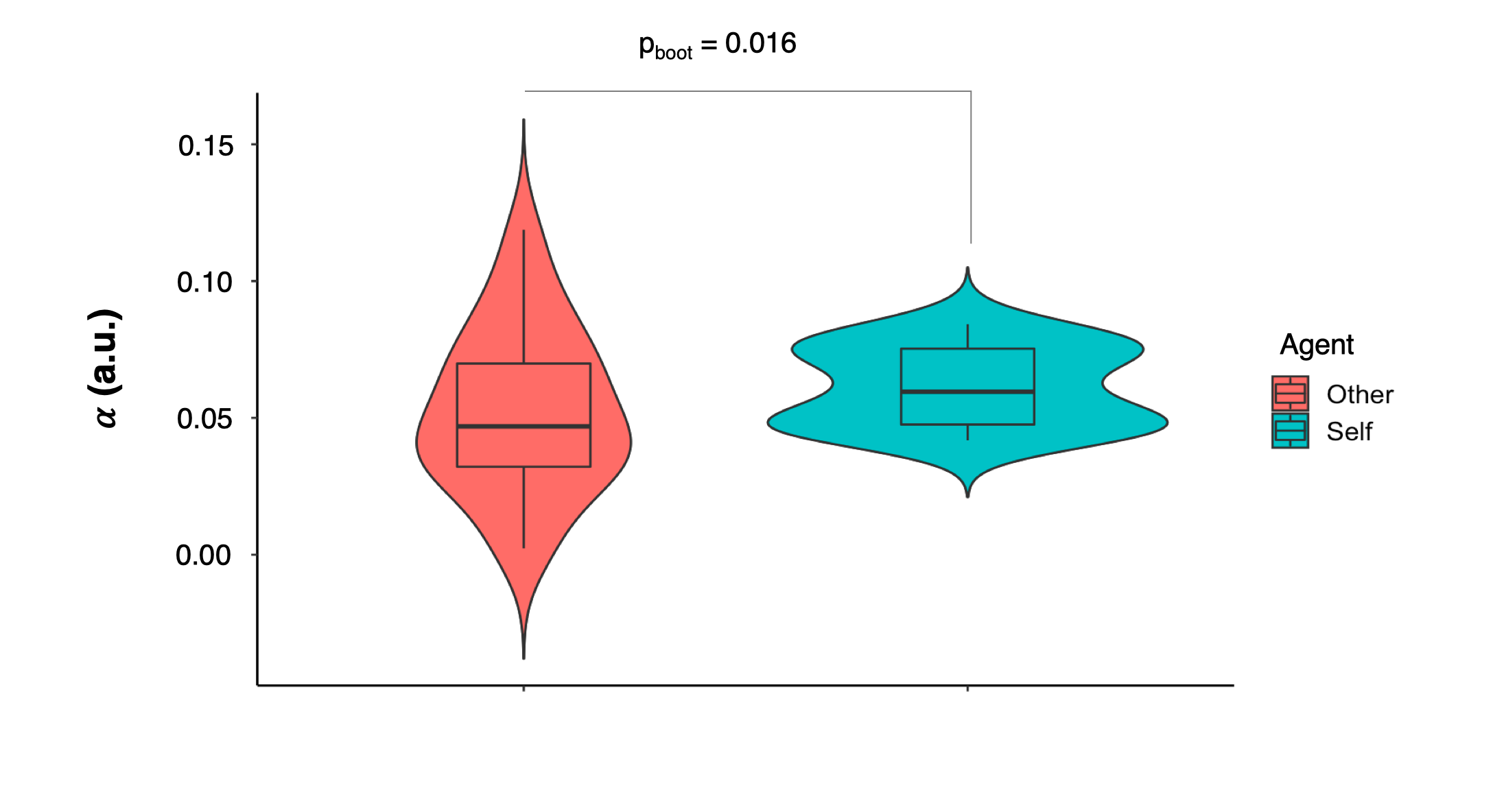

**Supplementary Figure S9. Effects of treatment on the inverse temperature parameter beta (β).** We examined the effects treatment on the inverse temperature parameter beta (β), using a linear mixed model where we specified random intercepts for participants. The effect of treatment was not significant. The violin plots show the distributions of β in each treatment level.

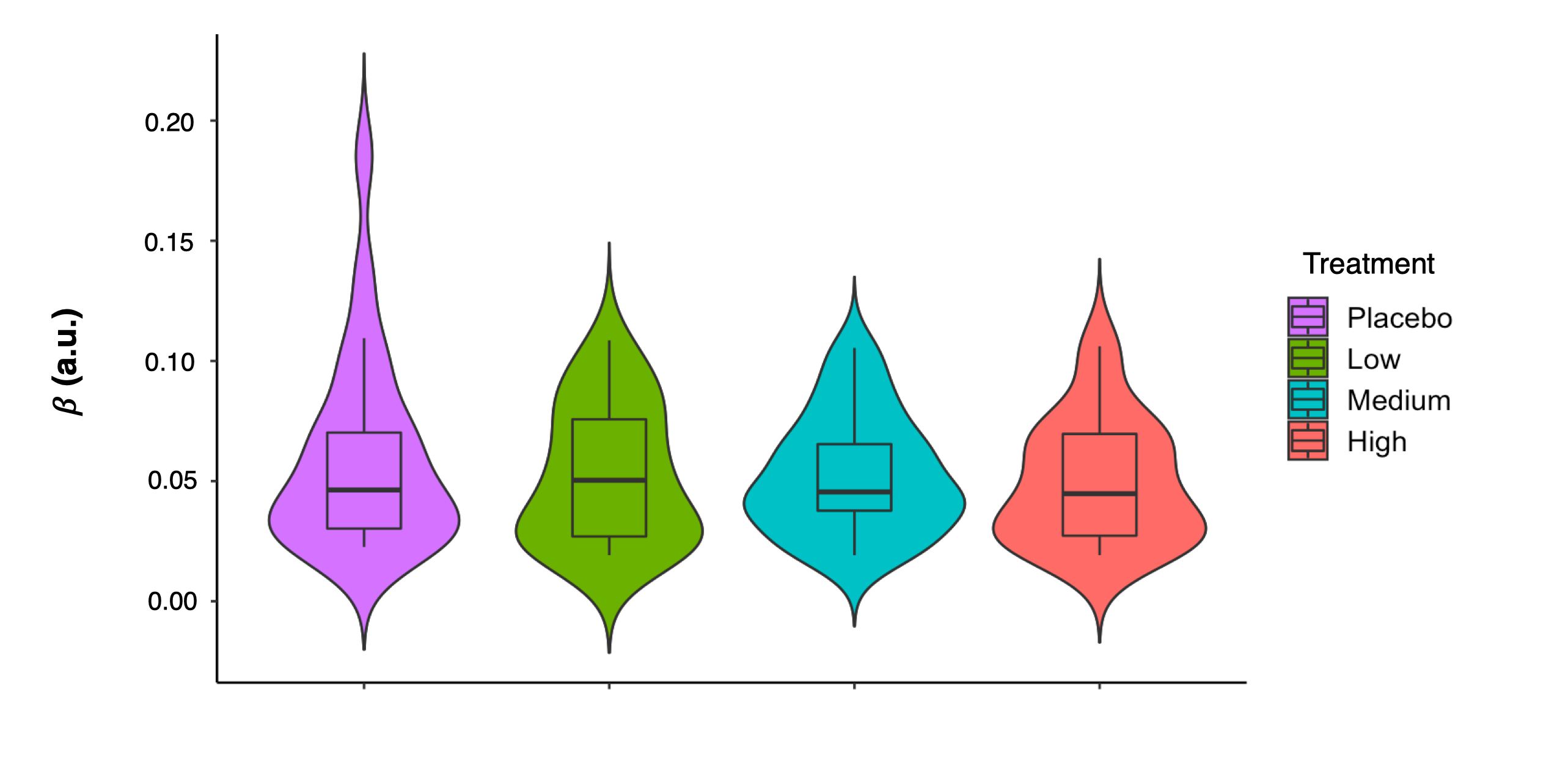

**Supplementary Figure S10. BOLD representations of prediction errors in the subgenual anterior cingulate, nucleus *accumbens* and midbrain.** Using data from the placebo level, first we investigated whether we could capture hypothesised encoding (BOLD representations) of prediction errors in the subgenual anterior cingulate (a), nucleus accumbens (b) and midbrain (c). Here, we provide violin plots of extracted parameter estimates capturing this encoding for each of these three regions-of-interest. In each region, we tested the effect of learning condition using linear mixed models, specifying random intercepts for participants. Significance was assessed using bootstrap (1000 samples).

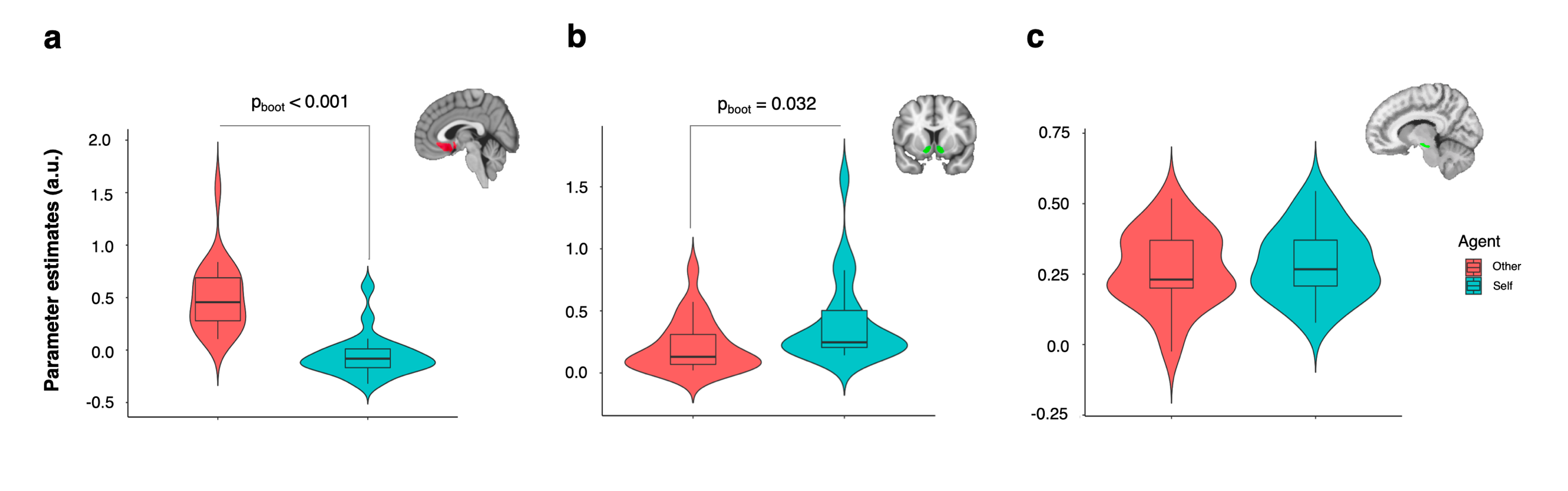

**Supplementary Figure S11. Correlations between encoding of prediction errors and learning performance.** We investigated whether encoding of prediction errors in each of our three regions-of-interest (a – subgenual anterior cingulate cortex; b – nucleus *accumbens*; c – midbrain) could capture inter-individual variability in the learning rates (α) of our participants during each of the two conditions. We calculated Pearson’s correlations with bootstrap (1000 samples). The lines represent the fit of a linear regression for each learning condition and the shadow the respective 95% confidence interval.

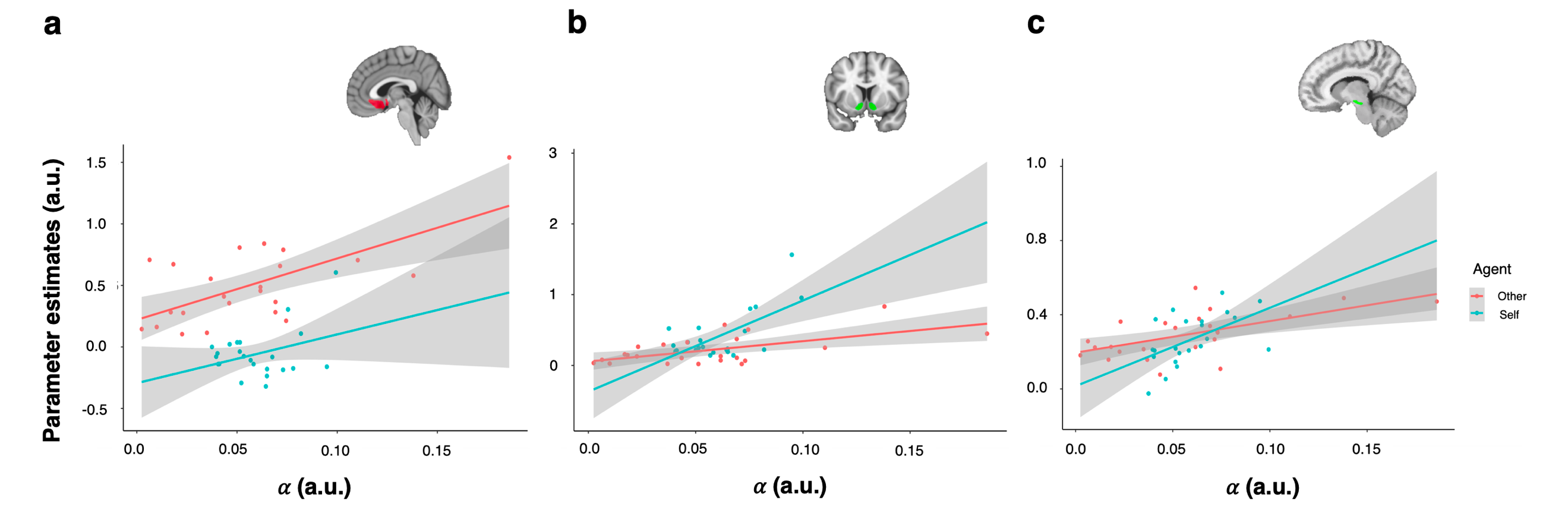

**Supplementary Figure S12. BOLD representations of prediction errors during self-oriented and prosocial learning (whole-brain analysis).** Here, we provide a representation of all clusters depicting a significant positive association with prediction errors during self-oriented (a) and prosocial (b) learning at the whole-brain level. We used data from the placebo level only and tested for positive and negative correlations with trial-by-trial trajectories of prediction errors, using directed One-sample T-tests. All clusters were significant at a corrected p_FWE_ < 0.05 threshold (cluster-forming threshold p<0.001, uncorrected). No clusters depicting negative correlations survived correction.

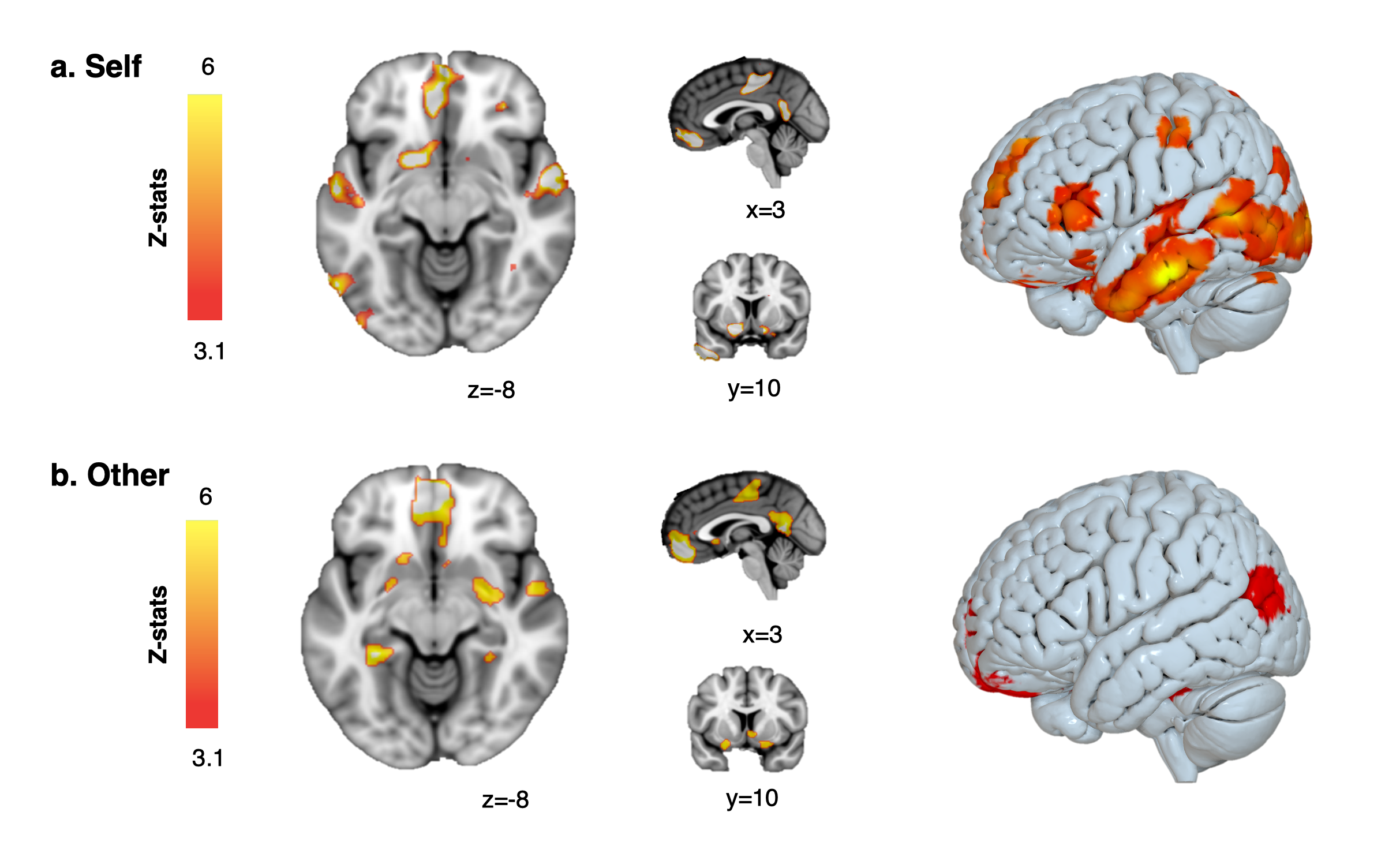

**Supplementary Figure S13. BOLD representations of expected value of the chosen actions (whole-brain analysis).** Here, we provide a representation of all clusters depicting a significant positive association with the expected value of the chosen actions during self-oriented (a) and prosocial (b) learning at the whole-brain level. We used data from the placebo level only and tested for positive and negative correlations with trial-by-trial trajectories of expected value of the chosen actions, using directed One-sample T-tests. All clusters were significant at a corrected p_FWE_ < 0.05 threshold (cluster-forming threshold p<0.001, uncorrected). No clusters depicting negative correlations survived correction.

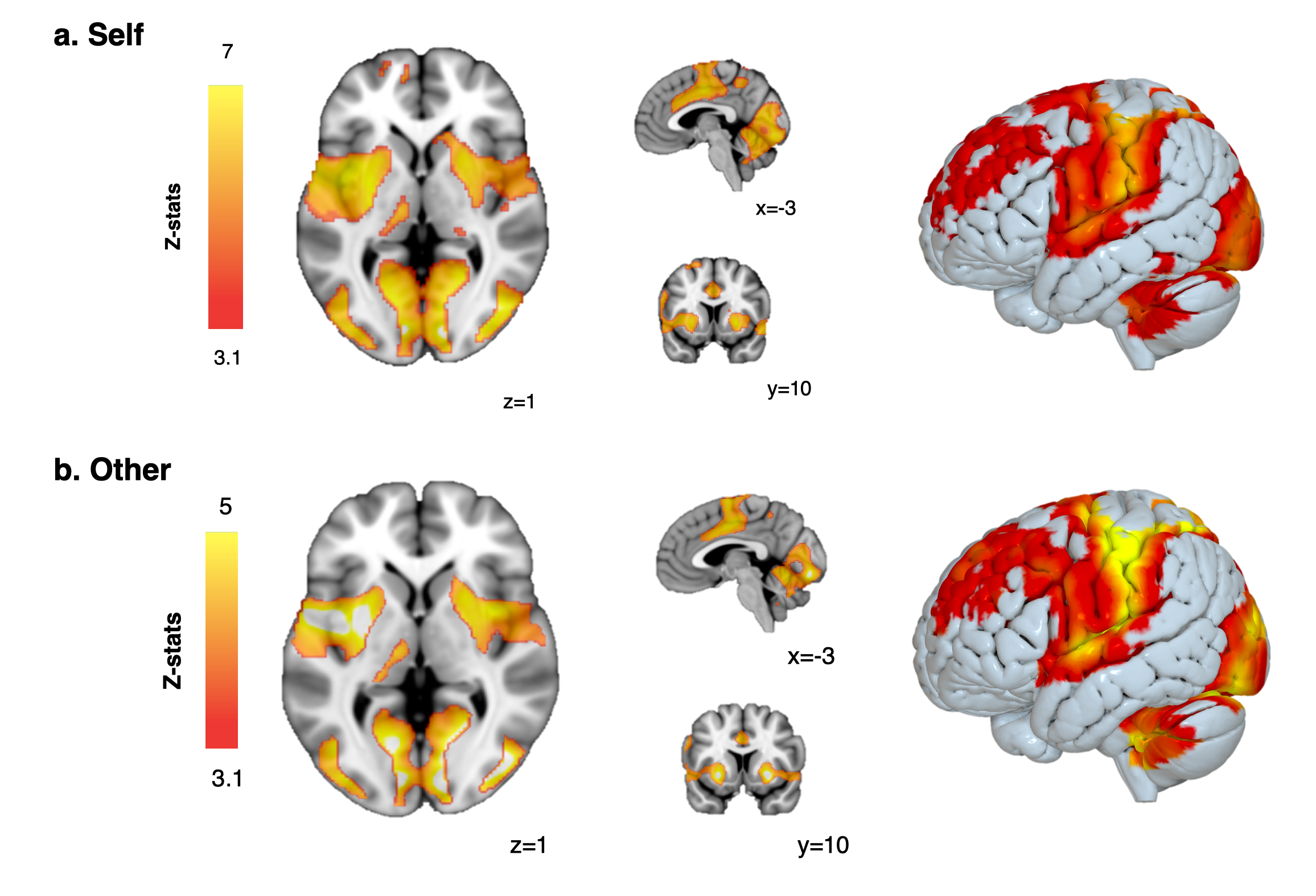

**Supplementary Table S3. Effects of learning condition, treatment and condition x treatment on encoding of prediction errors in the subgenual anterior cingulate cortex, nucleus accumbens and midbrain.** In this table, we present the results of the linear mixed models investigating the effects of condition, treatment and condition x treatment on encoding of prediction errors in the nucleus accumbens, subgenual anterior cingulate cortex and midbrain.

| **Region-of-interest** | **Main effect of treatment** | **Main effect of learning condition** | **Treatment x Learning condition interaction** |
| --- | --- | --- | --- |
| Nucleus accumbens | χ^2^ (3) = 0.185  p_boot_ = 0.972 | χ^2^ (1) = 18.803  p_boot_ < 0.001 | χ^2^ (3) = 0.408  p_boot_ = 0.945 |
| Subgenual anterior cingulate | χ^2^ (3) = 18.343  p_boot_ < 0.001 | χ^2^ (1) = 190.564  p_boot_ < 0.001 | χ^2^ (3) = 16.431  p_boot_ = 0.004 |
| Midbrain | χ^2^ (3) = 3.208  p_boot_ = 0.401 | χ^2^ (1) = 3.635  p_boot_ = 0.063 | χ^2^ (3) = 11.058  p_boot_ = 0.010 |

**Supplementary Table S4. Effect of learning condition x treatment on encoding of prediction errors in the subgenual anterior cingulate cortex and midbrain during self-oriented and prosocial learning (post hoc tests).** In this table, we present the results of the post hoc tests investigating the significant interaction learning condition x treatment we found for encoding of prediction errors in the subgenual anterior cingulate cortex (ACC) and midbrain. Statistical significance was set to p_adj_ < 0.05, after Holm-Bonferroni correction for multiple testing.

| **Region of interest** | **Condition** | **Stats** | **Low**  **vs Placebo** | **Medium vs Placebo** | **High**  **vs Placebo** | **Low**  **vs High** | **Low**  **vs Medium** | **High**  **vs Medium** |
| --- | --- | --- | --- | --- | --- | --- | --- | --- |
| Subgenual ACC | Self-oriented | t  p_adj_ | -0.755  1.000 | -0.538  1.000 | -1.002  1.000 | 0.247  1.000 | -0.217  1.000 | -0.464  1.000 |
|  | Prosocial | t  p_adj_ | 3.180  0.005 | -0.628  0.531 | -2.631  0.019 | 5.877  < 0.001 | 3.852  < 0.001 | -2.026  0.089 |
| Midbrain | Self-oriented | t  p_adj_ | -0.841  1.000 | -1.601  0.445 | 0.903  1.000 | -1.744  0.416 | 0.761  1.000 | 2.504  0.080 |
|  | Prosocial | t  p_adj_ | 2.566  0.067 | 1.128  0.783 | 0.381  0.912 | 2.185  0.152 | 1.438  0.610 | -0.747  0.912 |

**Supplementary Figure S14. Encoding of prediction errors in the functional coupling of the midbrain.** Using data from the placebo level, first we investigated whether we could capture encoding of prediction errors during self-oriented and prosocial learning in the functional coupling between the midbrain and subgenual anterior cingulate (a), and between the midbrain and the nucleus *accumbens* (b). Here, we provide violin plots of extracted psychophysiological interaction parameter estimates capturing this encoding. In each region, we tested for an effect of condition using linear mixed models, specifying random intercepts for participants. Significance was assessed using bootstrap (1000 samples).

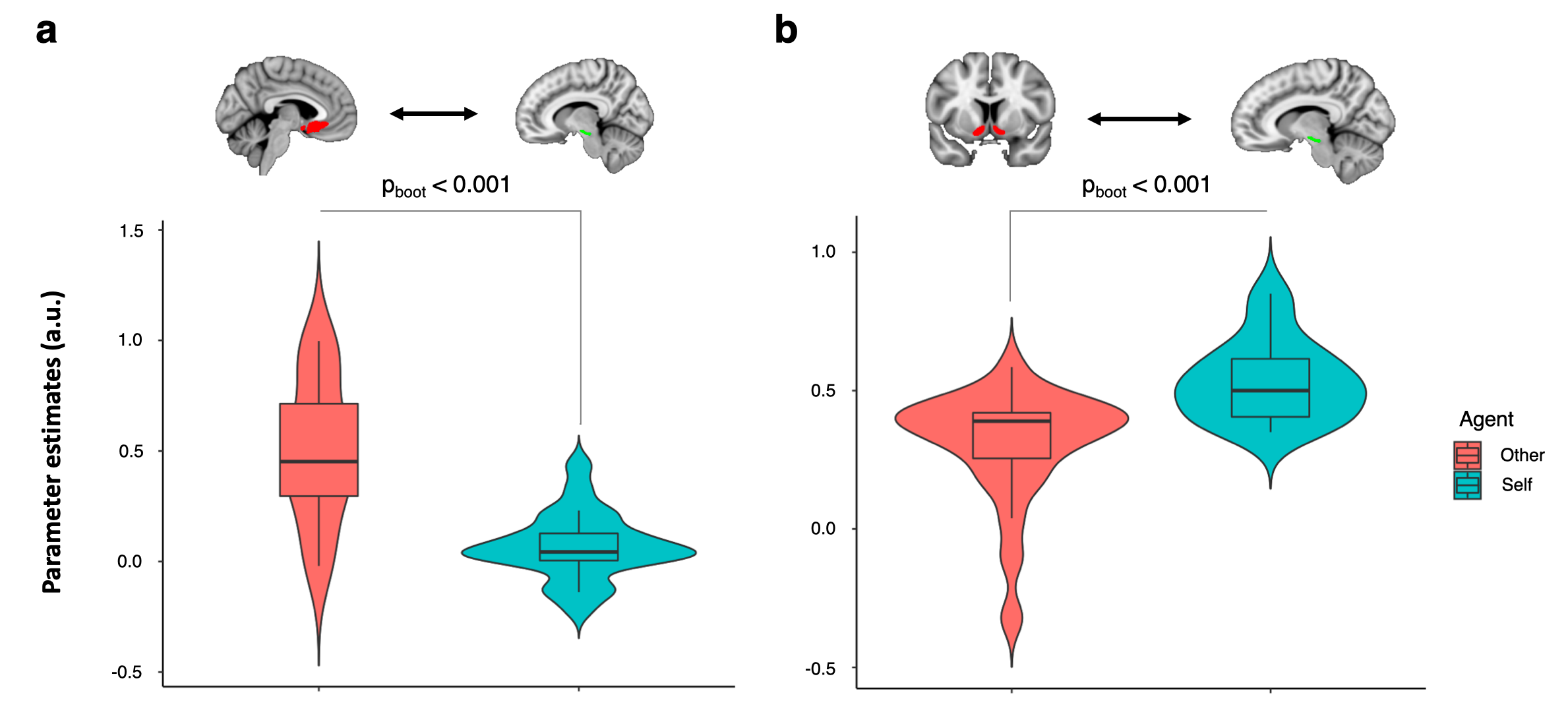

**Supplementary Figure S15. Correlations between encoding of prediction errors in the functional coupling of midbrain and performance during self-oriented and prosocial learning.** We investigated whether encoding of prediction errors in midbrain – subgenual anterior cingulate cortex functional coupling (a) could capture inter-individual variability in the learning rates (α) of our participants during each learning condition. We repeated the same analysis this time focusing on the midbrain – nucleus accumbens functional coupling (b). We calculated Pearson’s correlations with bootstrap (1000 samples). The lines represent the fit of a linear regression for each condition and the shadow the respective 95% confidence interval.

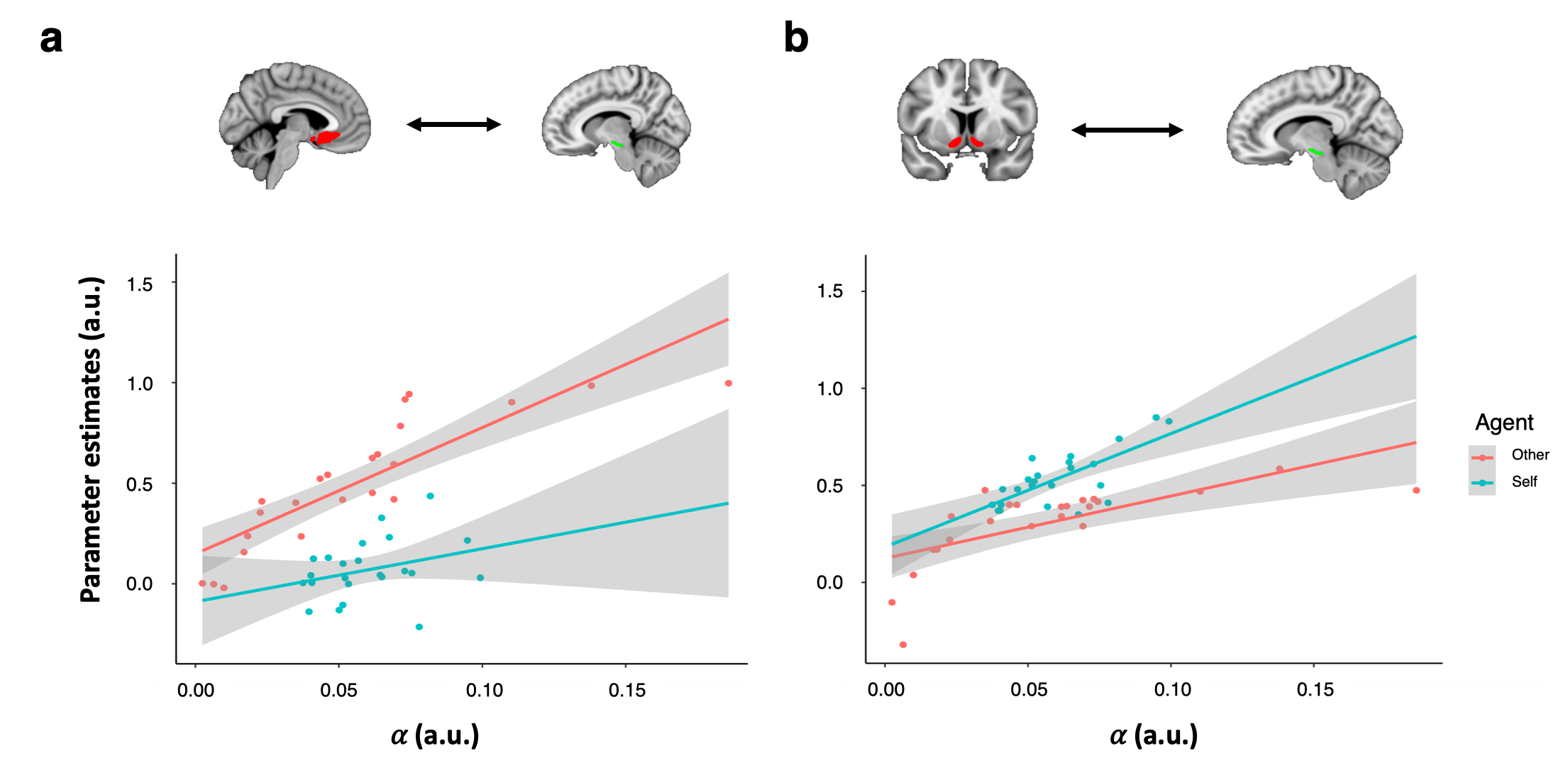

**Supplementary Table S5. Effects of condition, treatment and condition x treatment on encoding of prediction errors in the functional coupling between the midbrain and the subgenual anterior cingulate cortex, and between the midbrain and nucleus *accumbens*.** In this table, we present the results of the linear mixed models investigating the effects of learning condition, treatment and learning condition x treatment on encoding of prediction errors in the functional coupling between the midbrain and subgenual anterior cingulate cortex, and between the midbrain and nucleus accumbens.

| **Region-of-interest** | **Main effect of treatment** | **Main effect of learning condition** | **Treatment x learning condition interaction** |
| --- | --- | --- | --- |
| Midbrain – Subgenual anterior cingulate | χ^2^ (3) = 18.620  p_boot_ < 0.001 | χ^2^ (1) = 149.425  p_boot_ < 0.001 | χ^2^ (3) = 15.727  p_boot_ < 0.001 |
| Midbrain – nucleus accumbens | χ^2^ (3) = 2.343  p_boot_ = 0.525 | χ^2^(1)=109.904, p_boot_<0.001 | χ^2^ (3) = 1.028  p_boot_ = 0.810 |

**Supplementary Table S6. Effect of learning condition x treatment interaction on encoding of prediction errors in the functional coupling between the midbrain and subgenual anterior cingulate cortex.** In this table, we present the results of the post hoc tests investigating the significant interaction learning condition x treatment we found for encoding of prediction errors in the functional coupling between the midbrain and subgenual anterior cingulate cortex (ACC). Statistical significance was set to p_adj_ < 0.05, after Holm-Bonferroni correction for multiple testing.

| **Region of interest** | **Agent** | **Stats** | **Low**  **vs Placebo** | **Medium vs Placebo** | **High**  **vs Placebo** | **Low**  **vs High** | **Low**  **vs Medium** | **High**  **vs Medium** |
| --- | --- | --- | --- | --- | --- | --- | --- | --- |
| Midbrain  - Subgenual ACC | Self-oriented | t  p_adj_ | 0.127  1.000 | -0.680  1.000 | 0.011  1.000 | 0.116  1.000 | 0.807  1.000 | 0.691  1.000 |
|  | Prosocial | t  p_adj_ | 2.911  0.015 | -0.870  0.386 | -2.949  0.015 | 5.861  < 0.001 | 3.782  0.001 | -2.079  0.078 |

**Supplementary Figure S16. Associations between learning rates and the effective connectivity between the midbrain and subgenual anterior cingulate cortex during the prosocial blocks.** We investigated associations between the learning rates (α) of our participants during self-oriented and prosocial learning and the effective connectivity between the midbrain and subgenual anterior cingulate cortex during the prosocial blocks in the placebo condition. We followed the same approach we used in our main dynamic causal modelling analysis of treatment effects, but this time our second-level parametrical empirical bayes (PEB) design matrix for the second level analysis included three regressors: i) mean; ii) correlation with learning rates during self-oriented blocks (α_self-oriented_); iii) correlation with learning rates during prosocial blocks (α_prosocial_). Our second level PEB models (a – upper panel) included four competing models with all possible combinations of learning rate effects. M_3_ was the winning model with the highest posterior probability and the lowest free energy (a – lower panel). This model included effects for the α_prosocial_ regressor only. We investigated this effect further by looking at the expected estimates and posterior probabilities (P_p_) of each parameter of the reduced PEB model. In panel B, we provide a schematic diagram of these effects. Grey/black lines present the mean expected estimates. In green, we present the effect of α_prosocial_. Bold lines indicate strong evidence in favour of an expected estimate reliably different from 0 (Pp>0.90). sgACC – Subgenual anterior cingulate cortex; P_p_ – Posterior probability.

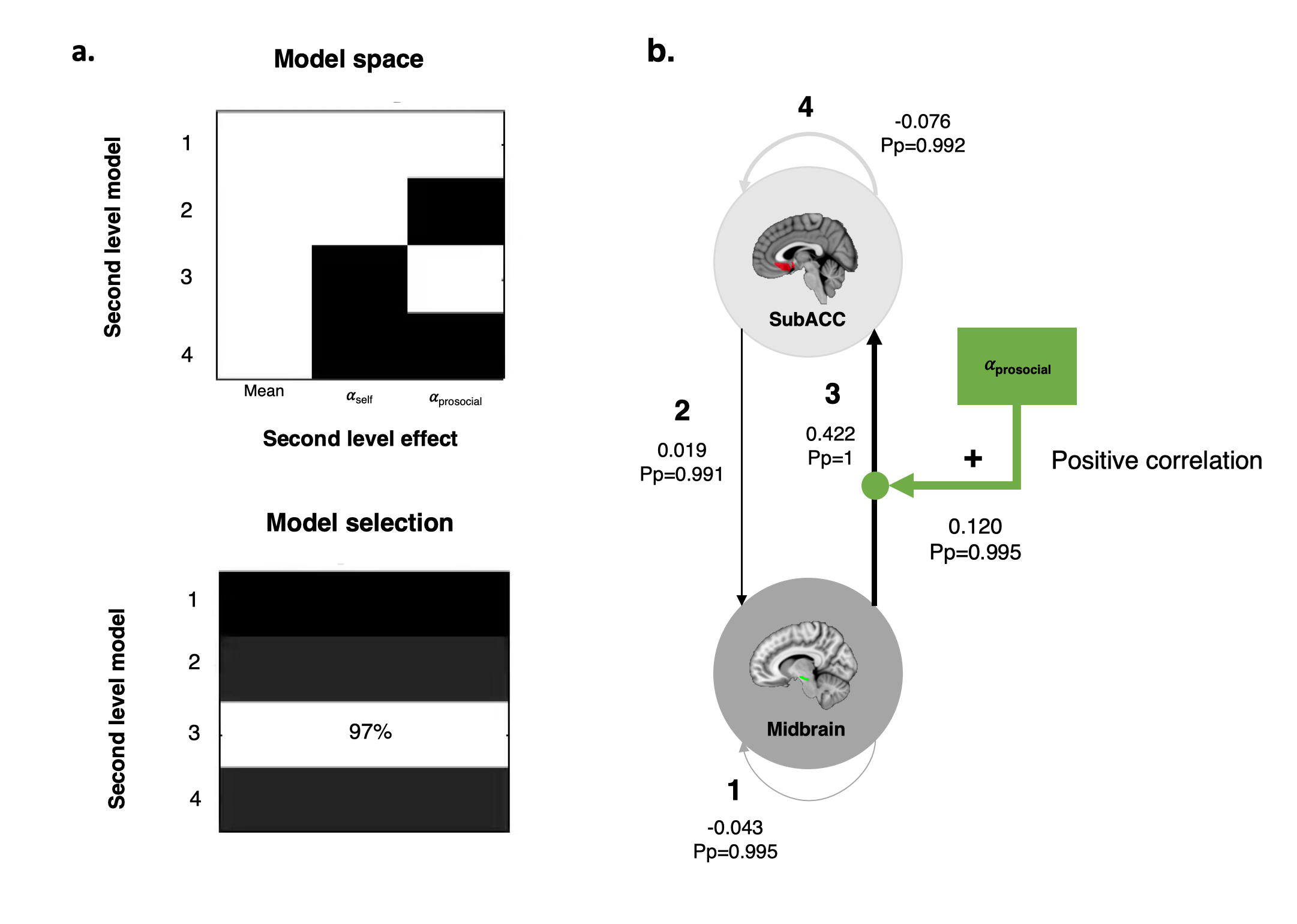
